# The pregnancy-associated protein glycodelin as a potential sex-specific target for resistance to immunotherapy in non-small cell lung cancer

**DOI:** 10.1101/2023.03.02.530822

**Authors:** Sarah Richtmann, Sebastian Marwitz, Thomas Muley, Hannu Koistinen, Petros Christopoulos, Michael Thomas, Daniel Kazdal, Michael Allgäuer, Hauke Winter, Torsten Goldmann, Michael Meister, Ursula Klingmüller, Marc A. Schneider

**Affiliations:** Translational Research Unit, Thoraxklinik at Heidelberg University Hospital, Heidelberg, Germany; Translational Lung Research Center Heidelberg (TLRC), Member of the German Center for Lung Research (DZL), Heidelberg, Germany; Division of Systems Biology of Signal Transduction, German Cancer Research Center (DKFZ), Heidelberg, Germany; Faculty of Biosciences, University of Heidelberg, Heidelberg, Germany; Histology, Research Center Borstel - Leibniz Lung Center, Borstel, Germany; Airway Research Center North (ARCN), Member of the German Center for Lung Research (DZL), Borstel, Germany; Department of Clinical Chemistry and Haematology, Faculty of Medicine, University of Helsinki and Helsinki University Hospital, Finland; Department of Thoracic Oncology, Thoraxklinik at Heidelberg University Hospital, Heidelberg, Germany; Institute of Pathology, Heidelberg University Hospital, Heidelberg, Germany; Department of Surgery, Thoraxklinik at Heidelberg University Hospital, Heidelberg, Germany

**Keywords:** NSCLC, Glycodelin, Immunotherapy, Tumor microenvironment, Glycosylation

## Abstract

Lung cancer has been shown to be targetable by novel immunotherapies which reactivate the immune system and enable tumor cell killing. However, treatment failure and resistance to these therapies is common. Consideration of sex as a factor influencing therapy resistance is still rare. We hypothesize that the success of the treatment is impaired by the presence of the immunosuppressive pregnancy-associated glycoprotein glycodelin that is expressed in patients with non-small-cell lung cancer (NSCLC). We demonstrate that the glycan pattern of NSCLC-derived glycodelin detected by a lectin-based enrichment assay highly resembles amniotic fluid-derived glycodelin A, which is known to have immunosuppressive properties. NSCLC-derived glycodelin interacts with immune cells *in vitro* and regulates the expression of genes associated with inflammatory and tumor microenvironment pathways. In tumor microarray samples of patients, high glycodelin staining in tumor areas results in an impaired overall survival of female patients. Moreover, glycodelin colocalizes to tumor infiltrating CD8+ T cells and pro-tumorigenic M2 macrophages. High serum concentrations of glycodelin prior to immunotherapy are associated with a poor progression-free survival (p < 0.001) of female patients receiving PD-(L)1 inhibitors. In summary, our findings suggest that glycodelin not only is a promising immunological biomarker for early identification of female patients that do not benefit from the costly immunotherapy, but also represents a promising immunotherapeutic target in NSCLC to improve therapeutic options in lung cancer.

**Background:** Immunotherapy is one of the major achievements in the last decade of lung cancer treatment. However, resistance to treatment is common und not well understood. Glycodelin is an immunosuppressive protein well described during the establishment of a pregnancy. We investigated its influence on immune cells and patients receiving immunotherapy with a focus on the sex of the patients.

**Translational relevance:** Our study examined that NSCLC-derived glycodelin shares similarities to amniotic fluid-derived glycodelin A and is predictive for a worse response to immunotherapy in female patients. Therefore, glycodelin might be a key player influencing a sex-specific response to immunotherapy in lung cancer.

## Introduction

Lung cancer is one of the most common and deadliest cancers worldwide, with an incidence of 2.2 million new cases per year and 1.8 million deaths. Immunotherapies have revolutionized the landscape of cancer treatment and have enabled long-term survival in patients with metastatic non-small cell lung cancer (NSCLC) (1). However, not all tumors that are positive for PD-L1 show immune infiltration, and some do not respond to anti-PD-1 therapies (2, 3). Consequently, additional markers will be needed that cover other immune checkpoint molecules expressed on tumor cells and surrounding cells in the tumor microenvironment.

In the 1980s several investigators identified a protein, which eventually was named as glycodelin, in human placenta, in amniotic fluid, the pregnancy decidua, and seminal plasma (4). Glycodelin is encoded by the *progesterone-associated endometrial protein* (*PAEP*) gene and a translated from 900 bp mRNA that shares sequence similarity with bovine β-lactoglobulin (5) and other lipocalins (6). Glycodelin has been proposed to be involved in immunosuppression, angiogenesis, and apoptosis signaling during the first trimester of pregnancy and to regulate fertilization and implantation (7–9) by immunomodulatory processes. Glycodelin A is found in amniotic fluid, in the secretory and decidualized endometrium and can also be detected in the serum of women who are pre-menopausal (7, 8, 10). The different functions of the glycodelin-glycoforms are based on the specific glycans at Asn28 and Asn63 (9). During pregnancy, Glycodelin A is the major form of glycodelin and acts as an immunomodulator on various levels. While the glycodelin has originally been found in reproductive tissues, it is also expressed in several cancers (11–13). In lung cancer, *PAEP* gene expression correlated with a significantly worse overall survival in female patients (13).

In the past years, immunotherapy represented a major improvement in the treatment of NSCLC. Since it plays an important role in the regulation of immunosuppressive pathways during the establishment of pregnancy in the female body, we characterized the interaction of glycodelin and the immune system in NSCLC and investigated a potential sex-specific influence of glycodelin on the success of immunotherapy.

## Material and methods

### Cell cultures

The cells were cultured under humidified conditions at 37 °C and 5 % CO_2_. The patient-derived fibroblast cell line 4950T-F and the LUSC cell line 2427T (14) were cultivated in DMEM/HAMs F12 (Thermo Fisher Scientific, Waltham, Massachusetts, US) with 10 % fetal bovine serum (FBS) and 1 x stable glutamine (both Thermo Fisher Scientific). For Jurkat, THP1, and KHYG-1 (all from ATCC), RPMI 1640 (Thermo Fisher Scientific) added with 10 % FBS was used. For KHYG-1, also 10 ng/ml IL-2 (PHC0026, ThermoFisher Scientific) was added. The patient derived primary cell lines 4950T and 170162T were cultivated in serum-free DMEM/HAMs F12 with epithelial airway growth factors (C-21160, PromoCell, Heidelberg, Germany) and ROCK inhibitor (Y-27632, Merck, Darmstadt, Germany).

### Transient gene knockdown by siRNA transfection

Cells were seeded in T75 culture flasks with a cell density of 1.3 x 10^6^ cells/flask and kept in the incubator at 37 °C and 5 % CO_2_ overnight. For one reaction, 15 µl of siRNA (1 µM, FlexiTube GeneSolution, Qiagen, Hilden, Germany, see Table S1) were diluted in 1.5 ml culture medium without supplements. 18 µl RNAiMax (Thermo Fisher Scientific) were added and the solution was incubated for 10-20 min at room temperature. The cell culture medium was exchanged to a final volume of 13.5 ml/flask and 1.5 ml siRNA solution were carefully dripped onto the cells, resulting in a final siRNA concentration of 1 nM. Treated cells were incubated for 72 h at 37 °C and 5 % CO_2_. Four siRNAs (#SI00039704, #SI00039711, #SI00039718, #SI03102659, all Qiagen) were used for glycodelin siRNA knockdown. As a control, AllStars Negative Control siRNA (#SI03650318, Qiagen) was used.

### Total RNA isolation and cDNA synthesis

For RNA isolation from cell lines, the RNeasy Mini Kit (Qiagen) was used. The complete procedure is described elsewhere (13).

### Quantitative real time polymerase chain reaction (qPCR)

*Real-time* quantitative PCR (qPCR) was performed in accordance with MIQE-guidelines (15) using a LightCycler^®^ 480 Real-Time PCR Instrument (Roche, Mannheim, Germany) in a 384-well plate format. Gene-specific primers and probes (Universal ProbeLibrary, Roche) were used in combination with qPCR Probe-MasterMix (SL-9802, Steinbrenner, Wiesenbach, Germany). 5 ng sscDNA were used for each amplification. The whole procedure is described elsewhere (13). The following forward and reverse primers and probes were used: ESD, 5’-TCAGTCTGCTTCAGAACATGG-3‘ and 5’-CCTTTAATATTGCAGCCACGA-3’, UPL50; RPS18: 5’-CTTCCACAGGAGGCCTACAC-3’ and 5’-CGCAAAATATGCTGGAACTTT-3’, UPL46; PAEP: 5’-CCTGTTTCTCTGCCTACAGGA-3’ and 5’-CGTCCTCCACCAGGACTCT- 3’, UPL77; CXCL10: 5’-GAAAGCAGTTAGCAAGGAAAGGT-3’ and 5’- GACATATACTCCATGTAGGGAAGTGA-3’, UPL34; NFKB: 5’-CCTGGAACCACGCCTCTA- 3’ and 5’-GGCTCATATGGTTTCCCATTTA-3’, UPL49; TNF: 5’- CAGCCTCTTCTCCTTCCTGAT-3’ and 5’-GCCAGAGGGCTGATTAGAGA-3’, UPL29; PDGFA: 5’-GATGAGGACCTTGGCTTGC-3’ and 5’-CCAGCCTCTCGATCACCTC-3’, UPL68; MMP9: 5’-GAACCAATCTCACCGACAGG-3’ and 5’-GCCACCCGAGTGTAACCATA-3’, UPL21; THBS1: 5’-GCAGGAAGACTATGACAA-3’ and 5’-CTGTCATCTGGAATTTTATCA-3’. RSPO3: 5’-GAAGCAGATTGGAGTATG-3’ and 5’-TCCACTTTTACATTTTGTG-3’. For THBS1 and RSPO3, primers and probe were designed by the assay design center from Sigma-Aldrich Oligoarchitect^TM^ since the UPL system was available until December 2020.

### Patient specimens

Patient specimens were collected at Thoraxklinik-Heidelberg and stored at Lung Biobank Heidelberg. All patients signed an informed consent. The local ethics committees of the Medical Faculty Heidelberg approved the use of biomaterial and data (S-270/2001 for biobanking of specimen, S-827/2020 for the glycodelin project).

The study was performed according to the principles set out in the WMA Declaration of Helsinki. Clinical parameters of the investigated patient cohorts are shown in Table 1.

### Tissue sample collection, characterization and preparation

Tissue samples and tissue multi arrays (TMA) from patients with NSCLC (see table 1) were provided by the Lung Biobank Heidelberg, a member of the accredited Tissue Bank of the National Center for Tumor Diseases (NCT) Heidelberg, the BioMaterialBank Heidelberg, and the Biobank platform of the German Center for Lung Research (DZL). All histopathological diagnoses were made according to the 2015 WHO classification for lung cancer by at least two experienced pathologists. Tumor stage was designated according to the 7th edition of the UICC tumor, node, and metastasis. Before TMA construction, a hematoxylin and eosin (H&E)-stained slide of each block was analyzed to select tumor-containing regions. A TMA machine (AlphaMetrix Biotech, Roedermark, Germany) was used to extract two tandem 1.0-mm cylindrical core samples from each tissue donor block.

### Multiplex Immunofluorescence

The multiplex immunofluorescence was performed with two different panels including either T cell or macrophage related markers. The following antibodies were used: glycodelin 1:1000 (N-20, Santa Cruz Biotechnology, Dallas, TX), CD68 1:100 (PG-M1, Agilent, Santa Clara, CA), panCK 1:300 (AE1/AE3, Zytomed, Berlin, Germany), iNos 1:300 (SP126, Abcam, Camebridge, UK), CD8 1:100 (SP16, Abcam), CD4 1:100 (EP204, Cell Signaling Technology, Danvers, MA), Granzyme B 1:300 (D6e9W, Cell Signaling Technology). The staining was performed in alternating steps, starting with antigen retrieval in AR6 buffer (Akoya Biosciences, Marlborough, MA) by heating in microwave oven for one min at 1250 W, followed by 10 min at 125 W. In the first round, endogenous peroxidase activity was removed by incubation with 3 % peroxide for 10 min. The slides were washed once in 0.1%Tween-20/Tris (pH 7.6) wash buffer and blocked in Akoya blocking buffer for 10 min at room temperature. Primary antibody was added in Renaissance Background Reducing Diluent (Zytomed) for 45 min at room temperature in the dark with gentle shaking. The slides were washed and anti-mouse/anti-rabbit HRP polymer (Akoya Biosciences) was added for 10 min at room temperature. Finally, the respective Tyramide-signal amplification (TSA) reaction with OPAL fluorochromes was performed with each OPAL-TSA conjugate diluted 1:150 in TSA plus reaction buffer with 10 min of enzyme reaction time. The next staining step was again initiated by antigen retrieval. panCK staining was performed by using an OPAL TSA-DIG antibody (Akoya Biosciences) in combination with an OPAL 780 fluorophore conjugated anti-DIG antibody. Cell nuclei were stained with spectral DAPI in PBS for 5 min and the slides were mounted with Hard-set Vectashield mounting medium (Vector Laboratories, Burlingame, CA). Mounted slides were allowed to harden prior to scanning.

For image acquisition, the mIF stained slides were scanned on a Vectra Polaris (Akoya Biosciences) as a.qptiff file at 0.5 µm pixel resolution using the 20× objective with saturation protection as a whole-slide overview. TMA cores were annotated using the TMA function of the Phenochart software (Akoya Biosciences) by setting a grid with 1.2Lmm punch diameter.

InForm V.2.4.1 and the PhenoptR R package were used for subsequent image analysis. Slides stained with the same panel were also included in the same inForm project. Multiple representative images representing the observed variability for each protein marker were selected for training purposes within inForm software. User-guided training for tissue segmentation or phenotyping was performed. When the test analysis resulted in satisfying classification regarding tissue segmentation, cell segmentation, and phenotyping, the algorithm was used for batch analysis among all images. Consistently misclassified images and results were omitted rigorously.

Machine learning-based tissue segmentation was applied using inForm software with the three different tissue categories ‘Tumor’, ‘Stroma’ and ‘Other’. User-annotated training regions for tumor identification included regions with a low expression of panCK or glycodelin and different histological subtypes to cover tissue heterogeneity. Overall tissue segmentation accuracy among the different staining panels was at least 95%. The cell segmentation algorithm from the inForm software V.2.4.1 was used and improved manually. Machine learning-based classification and counting of cellular phenotypes was performed by the use of inForm software on the protein markers used in the project. Selection of representative cellular phenotypes was done by manual annotation. For overall survival analyses, patients with incomplete surgical resection and no detectable glycodelin staining were excluded. The patient cohort is characterized in Table 1.

### Immunoblot

Samples were prepared with hot reducing SDS sample buffer containing 2-mercaptoethanol (Merck, Darmstadt, Germany) and loaded onto a 15 % SDS polyacrylamide gel to separate the proteins according to their size. The proteins were transferred onto a nitrocellulose membrane (pore size 0.2 µm) and transfer was confirmed by ponceau red staining. The membrane was blocked with 5 % (w/v) skimmed milk in 0.1 % (v/v) Tween20/PBS for 1 h at room temperature. Primary antibodies were added in skimmed milk buffer overnight at 4 °C. Glycodelin was detected using the polyclonal goat anti-glycodelin antibody 1:300 (N-20, Santa Cruz Biotechnology). Mouse anti-beta-actin 1:10,000 (A5441, Merck, Darmstadt, Germany) was used as loading control. The membrane was washed 4 times for 5 min with 1 x PBS/Tween20 0.1 % (v/v) and the HRP conjugated secondary antibodies (A5420 and A4416, Merck) were added 1:5000 (goat) and 1:10000 (mouse) for 1 h at room temperature. The membrane was washed, ECL substrate was added to initiate a chemiluminescent reaction and the signal was either detected by film exposure or by chemiluminescence imaging system ECL ChemoStar Imager HR 9.0 (Intas, Göttingen, Germany).

### Lectin-based pull-down assay

A set of eight different lectins (BK-1000, BK-3000, BK-2100, BIOZOL, Eching, Germany) was used for glycan binding studies. Biotinylated lectins were incubated with streptavidin-coupled magnetic beads (Thermo Fisher Scientific) for 45 min on the overhead shaker. Cell culture supernatants from 4950T or 170162T cells were concentrated 10x using Amicon centrifugal filtration units with 100 kDa cutoff filter (Merck). Concentrated supernatant or glycodelin A isolated from amniotic fluid was added to the lectin coupled beads and incubated overnight on an overhead shaker at 4 °C. Supernatant was collected as control and the beads were thoroughly washed 3 x with TBS/Tween 0.2% after which bound proteins were eluted by adding hot reducing SDS sample buffer containing 2-mercaptoethanol (Merck) and boiling for 5 min at 99 °C. Samples were analyzed *via* immunoblot. For each lectin, signal in control supernatant (flowthrough, FT) was compared to the signal of bound protein and scored according to the relative signal. Binding was considered was considered as strong (>70 % of the signal was detected in the bound fraction compared to the FT), moderate (40-69 % of the signal in bound fraction), weak (10-39 %), or no detectable binding (0-9 %).

### Measuring of glycodelin and hormones

Glycodelin levels in serum were measured using a glycodelin ELISA from Bioserv (BS-30-20, Bioserv Diagnostics, Berlin, Germany) according to the manufacturer’s instructions. Patient cohort is described in Table 1. Glycodelin concentrations in cell culture supernatants were measured using in house immunoassay (16).

The hormones were measured by the clinical diagnostics of the Heidelberg University Hospital. For human chorionic gonadotropin (hCG), progesterone and testosterone the ADVIA Centaur XPT system was used (Siemens Healthcare Diagnostics, Eschborn, Germany). Estrogen concentration was determined using Cobas E411 (Roche Diagnostics, Mannheim, Germany).

### *In vitro* binding assays and antibody blocking

The cell line 4950T secretes glycodelin into the culture supernatant in high amounts compared to other cell lines (average of 100 ng/ml for confluent cells per day). For cell-based assays, cell culture supernatant was concentrated using Amicon filter columns with either 30 kDa or 100 kDa cutoff (Merck), retaining glycodelin in the concentrated part. For time course binding assays, the concentrated supernatant was diluted to the initial concentration with cell culture medium without FBS. For binding assays with deglycosylated glycodelin, the condensed supernatant was first incubated with PNGase F (Promega, Walldorf, Germany), heat-inactivated, and then diluted to the initial concentration. Deglycosylation was controlled by immunoblot. For blocking assays, unconcentrated cell culture supernatants were incubated with a rabbit monoclonal anti-glycodelin antibody (clone 12C9, kindly provided by ROCHE, Penzberg, Germany) for 1 h at room temperature prior to incubation with immune cells. 5 µg of normal rabbit IgG (source) was used as control. After treatment, cells were washed three times with PBS and boiled for 5 min at 99°C in hot reducing SDS-buffer containing 2-mercaptoethanol (Merck).

### Affymetrix gene expression analysis

Control and knockdown 4950T cell culture supernatants were concentrated using an Amicon filter column with a 30 kDa cutoff and glycodelin levels were measured *via* ELISA. The supernatants were diluted to final glycodelin concentrations of 200 ng/ml (control) and 60 ng/ml (knockdown) in RPMI without FBS (+10 ng/ml IL-2 for KHYG-1 treatment). The immune cell lines Jurkat, THP1, and KHYG-1 were washed and 5 x 10^5^ cells were treated with 1 ml of concentrated supernatant for 3, 8, or 24 h. Cells were collected, washed with PBS and RNA was isolated as described above.

For Affymetrix gene chip analysis, total RNA was processed following the instructions of GeneChip™ 3’ IVT PLUS Reagent Kit User Guide (Manual Target Preparation for GeneChip™ 3’ Expression Arrays). Chips covering the Human Genome Array Type: HG-U133_Plus_2 were prepared and used to assess the gene expression. Data were analyzed using Transcriptome Analysis Console and Ingenuity Pathway Analysis Software (Qiagen). Genes with a fold change expression of < -2 or > 2 and a p-value < 0.05 were included in subsequent analyses. Canonical pathway analysis and networks were generated using QIAGEN IPA. The microarray data were deposited into the NCBI GEO database (accession number GSE222124).

### Statistical analyses

Analysis of survival data was performed using Kaplan-Meier estimator and Cox proportional hazards model and statistically analyzed under REMARK criteria using SPSS 29.0. Progression-free survival (PFS) time is calculated from the date of immunotherapy until the day of diagnosed tumor progression or death. The cut-off between high and low concentrations was identified by CutOff Finder version 2.1 (17). For glycodelin serum analyses, the median was the optimal cut-off. Univariate significance between the groups was examined by the log-rank test. Multivariate survival analysis was performed using the Cox proportional hazards model. Mann-Whitney U test was used to investigate significant differences between non-parametric datasets (patient related data). The Spearman ranked correlation coefficient test was performed for correlation analyses. Paired t-test was applied for *in vitro* experiments with at least three biological replicates. A p-value of less than 0.05 was considered significant. Visualization of the data was made by GraphPad Prism 5 (GraphPad Software, San Diego, CA).

## Results

### Comparison of the glycosylation pattern of NSCLC-derived glycodelin and immunosuppressive glycodelin A

The immunosuppressive function of glycodelin A is primarily mediated by its high sialylation degree and a distinct glycosylation pattern (18). In order to characterize the glycosylation pattern of NSCLC-derived glycodelin, a lectin based pull-down assay was performed investigating eight different binding specificities.

For these experiments, two patient-derived NSCLC cell lines 4950T and 170162T (Supplemental Fig. S1) were used. Both cell lines express the lung adenocarcinoma markers thyroid transcription factor 1 (TTF1) and cytokeratin 7 (CK7) and secrete high amounts of glycodelin. As outlined in Fig. 1A, the supernatants of the NSCLC cells were concentrated and incubated with different lectins and the binding of glycodelin was assessed by immunoblotting. The successful concentration of the cell culture supernatants through a centrifugal filter with a mass cutoff of 100 kDa was confirmed by the presence of glycodelin in the supernatant concentrates (Fig. 1B). The binding of glycodelin present in the supernatants from 4950T cells (SN1) and 170162T cells (SN2) to specific lectins was compared to the binding of glycodelin AF, isolated from amniotic fluid, that was used as a control glycodelin glycoform with a known immunosuppressive function (Fig. 1C).

**Figure 1:**
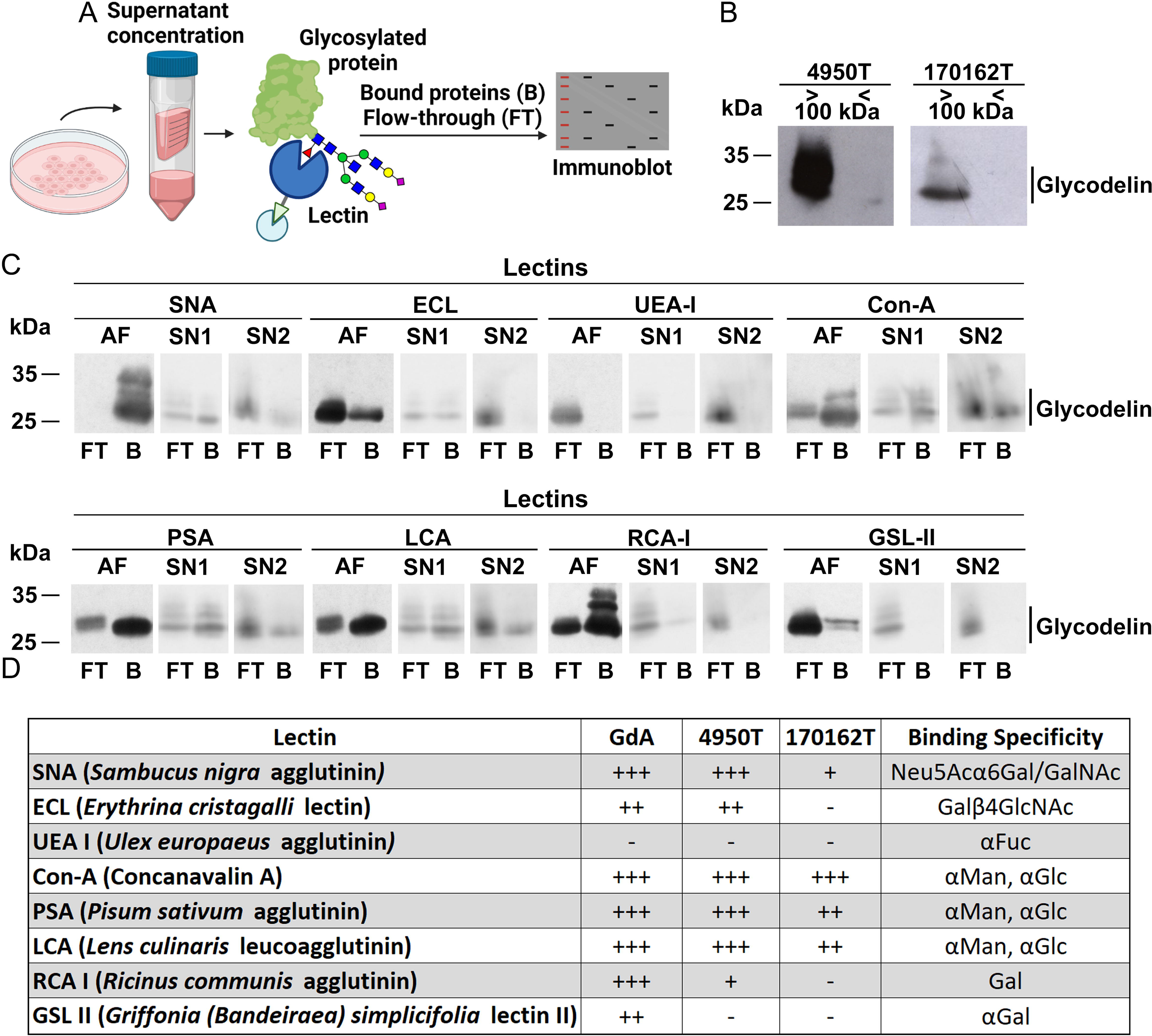
Comparison of the glycosylation pattern of NSCLC-derived glycodelin and glycodelin A. A) Schematic description of the experimental setup. Supernatant from cell lines 4950T and 170162T were collected, condensed, incubated with different lectins and analysed by immunoblot. B) Glycodelin in concentrate and flowtrough after concentration with a 100 kDa protein column. C) Binding capacities of glycodelin from amniotic fluid (AF), supernatant of 4950T (SN1) and 170162T (SN2) to eight different lectins. D) Evaluation of signal intensities from immunoblots shown in B). Binding specificity is based on the manufacturer’s (Vector Laboratories) information. AF = glycodelin from amniotic fluid, SN1 = 4950T supernatant, SN2 = 170162T supernatant, FT = flowthrough, B = bound to lectin, GdA = glycodelin A. +++ strong binding, ++ moderate binding, + weak binding, - no binding.

The results of the immunoblotting analysis (Fig. 1C and D) revealed that glycodelin present in the NSCLC cell line supernatants and immunosuppressive glycodelin A from amniotic fluid share many features. All examined glycodelins were bound by *Sambucus Nigra Agglutinin* (SNA) which is specific for sialic acid, an ubiquitous carbohydrate residue in glycodelin A important for its immunosuppressive function (19). However, 170162T derived glycodelin lacked the binding to *Erythrina cristagalli* lectin (ECL), while otherwise the glycodelins from 170162T and 4950T cell supernatants behaved similar in lectin binding assays. Glycodelin from all three sources were bound by Concanavalin-A (Con-A), *Pisum sativum* agglutinin (PSA), and *Lens culinaris* agglutinin (LCA) and thus seemed to share the typical sugar backbone consisting of mannose and glucose (20).

### Binding of NSCLC-derived glycodelin to immune cells *in vitro*

After revealing the structural similarities of the glycans of glycodelin A and NSCLC-derived glycodelin, we investigated the capability of 4950T-derived glycodelin to bind to immune cells. Binding of glycodelin to immortalized leukocyte cell lines Jurkat (T lymphocytes), THP1 (monocyte-derived), and KHYG-1 (natural killer cells) was analyzed by immunoblotting. Time course experiments revealed rapid binding kinetics of glycodelin to the three immune cell lines (Fig. 2A). We confirmed that the immune cells by themselves did not express glycodelin and that glycodelin was not detectable in the culturing media (Supplemental Fig. S2A).

**Fig. 2.**
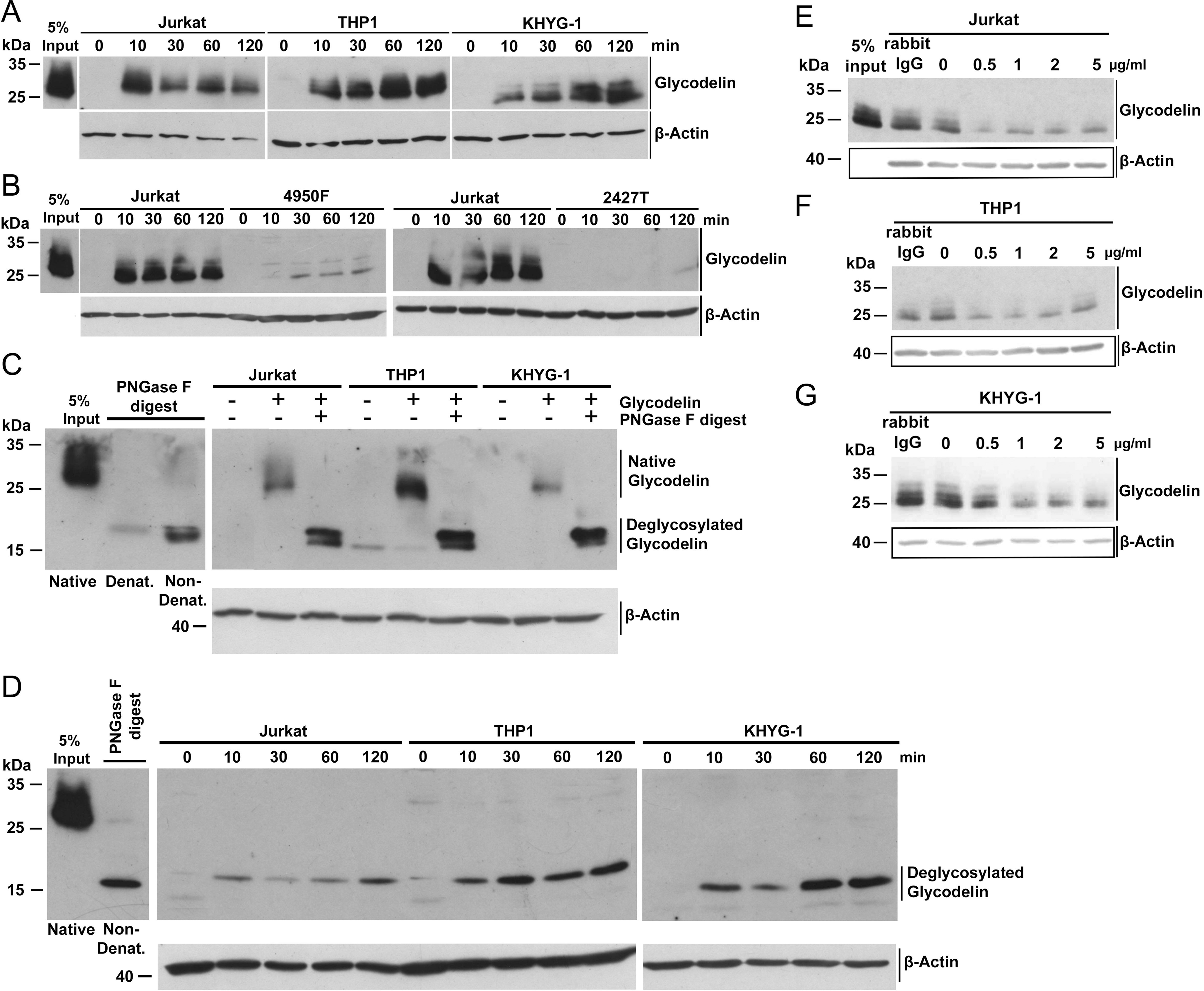
Binding of NSCLC-derived glycodelin to immune cells. A) The three immune cell lines Jurkat, THP1 and KHYG-1 were incubated with glycodelin-containing supernatants from 4950T cells for the indicated time. Cells were washed with PBS to remove unbound glycodelin and lysed for immunoblot. B) A fibroblast cell line 4950F as well as a lung squamous cell carcinoma cell line 2427T were incubated with glycodelin-containing supernatants from 4950T for the indicated time. C) Glycodelin from 4950T supernatant was deglycosylated by PNGase F digestion. Subsequently, the three cell lines Jurkat, THP1 and KHYG-1 were incubated with the undigested and digested supernatants. D) Time course of binding of deglycosylated glycodelin to the three investigated cell lines. Glycodelin was deglycosylated by digesting with PNGase F. Beta-actin was used as loading control. E-G) Supernatants of 4950T cells were preincubated with a monoclonal anti-glycodelin antibody for 1 h with the indicated concentrations. Afterwards, Jurkat (D), THP1 (E) and KHYG-1 (F) were cultivated in these supernatants for 24 h, washed and lysed for immunoblots. A normal rabbit IgG was used as negative control. Beta-actin was used as loading control.

The incubation of tumor-derived fibroblasts or of a lung squamous cell line with the supernatants of 4950T showed only poor or no binding of glycodelin supporting that binding of glycodelin is specific for immune cells (Fig. 2B). A pretreatment of glycodelin-containing supernatants with PNGase F to remove the glycosylation revealed that the binding of deglycosylated glycodelin to the immune cells was at least as strong and as rapid as the binding of the glycosylated form (Fig. 2C and D). The application of increasing anti-glycodelin antibody concentrations resulted in a strong inhibition of glycodelin binding (Fig. 2E-G) pointing a translational application.

### Impact of NSCLC derived glycodelin on gene expression in monocytic and natural killer cells

Numerous studies have investigated the highly pleiotropic effects and the modulation of the functionalities of different leukocytes through the interaction with glycodelin (19–21). To delineate the functional impact of NSCLC-derived glycodelin on immune cells, a gene knockdown was performed in 4950T cells targeting the glycodelin gene. The supernatants of knockdown cells were utilized to stimulate immune cells and the resulting alterations in gene expression was assessed by gene expression chip analysis (Fig. 3A). The knockdown efficiency increased with higher siRNA concentrations (Fig. 3B) but also led to a reduced cell viability (Supplemental Fig. S3A). The leucocyte cell lines THP1, Jurkat and KHYG-1 were cultivated with concentrated supernatants from 4950T cells treated with a scrambled siRNA (control) or 1 nm *PAEP* siRNA (Supplemental Fig. S3B). The leukocytes were incubated for 3, 8, and 24 h with the supernatants of 4950T cells to cover possible early and late gene expression responses. Of note, upon cultivation in the medium of 4950T cells, the viability of KHYG-1 was generally reduced to 50 to 60% (Supplemental Fig. S3C). For THP1 cells, up to 138 differentially expressed genes were observed at the three treatment time points, whereas for KHYG-1 cells, only at the 24 h time point 58 regulated genes were identified (Supplemental Table S1). For Jurkat cells, none of the treatments with glycodelin containing 4950T media resulted in a change of gene expression compared to media from glycodelin knockdown cells. Several inflammation related genes were upregulated genes in THP1. These included the tumor necrosis factor (*TNF)*, the C-X-C motif chemokine ligand 10 (*CXCL10)*, the chemokine (C-C motif) ligand 4 (*CCL4)*, and the intercellular adhesion molecule (*ICAM1*) (Supplemental Fig. S3D, Supplemental Table S1). In contrast, the gene thrombospondin 1 (*THBS1*) was downregulated in most of the analyzed samples. In KHYG-1 cells, similar to the findings in THP1 cells, genes encoding inflammation related factors such as *TNF*, *CCL4*, and interferon gamma (*IFNG)* as well as cell-cycle induction associated genes Cyclin E2 (*CCNE2*), Cell division control protein 6 homolog (*CDC6*), or the transcription factor *E2F8* were differentially expressed after glycodelin treatment.

**Fig. 3:**
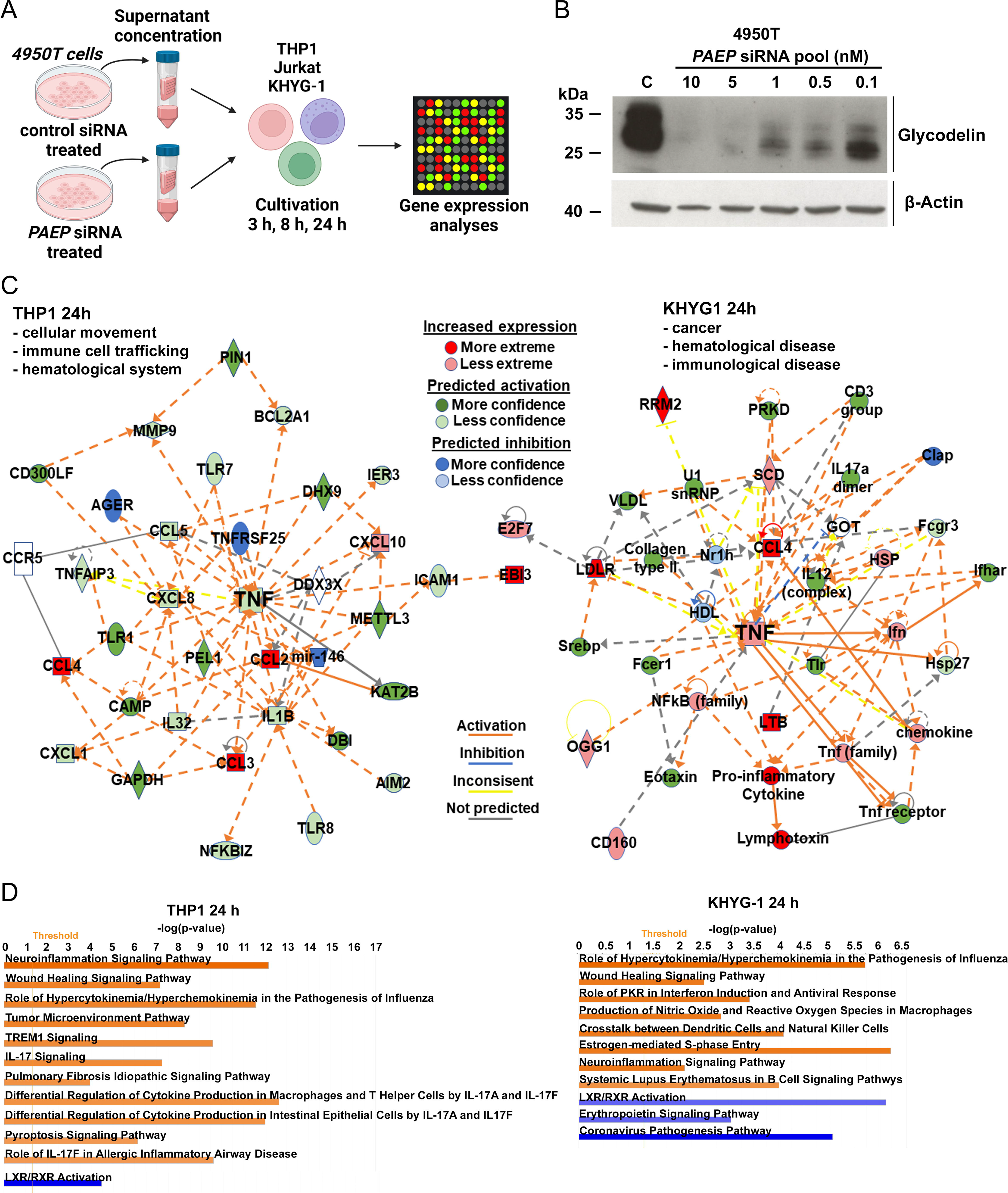
Effects of cultivation of THP1 and KHYG-1 cells with glycodelin-containing supernatants. A) Schematic overview of experimental setup. Supernatants of 4950T cells treated with scrambled siRNA or PAEP siRNAs were used for THP1 cell and KHYG-1 cell incubation. Changes in gene expression were measured by gene expression chip analyses. B) 4950T cells were treated with a PAEP siRNA pool for 72 h with the indicated concentrations. Knockdown efficiency was controlled by immunoblotting. Beta-actin was examined as loading control. C) Ingenuity pathway analyses of THP1 and KHYG-1 cells after 24 h of treatment with high and low glycodelin-containing supernatants. D) Transcriptome Analysis of the 24 h treatment of THP1 cells and KHYG-1 cells. Upregulated pathways are highlighted in orange, downregulated pathways in blue.

A transcriptional network analysis revealed that the TNF is at the center of the network in THP1 cells, along with additional immune response related genes (Fig. 3C, left). The incubation of THP1 cells with medium of 4950T containing high amounts of glycodelin for 24 h resulted in gene expression changes of pathways that are important for cellular movement, immune cell trafficking, and hematological system development and function. The respective analysis of glycodelin treated KHYG-1 cells revealed changes in a major transcriptional interaction network that is important in cancer, hematological, and immunological disease (Fig. 3C, right), while *TNF* again was a central node.

Ingenuity Pathway Analyses (IPA) were performed to decipher the effect of glycodelin on signaling pathways in the glycodelin stimulated immune cells (Fig. 3D). In THP1 cells, expression of inflammatory pathway genes was deregulated such as the Triggering receptor expressed on myeloid cells 1 (TREM1) signaling pathway or neuroinflammation signaling pathway. In addition, signaling pathways associated with the tumor microenvironment were affected in all samples and the pathway regarding the regulation of epithelial mesenchymal transition by growth factors was already influenced after 3 and 8 h of the treatment, respectively. In KHYG-1 cells, similar inflammatory pathways were influenced by the treatment with medium containing high glycodelin concentrations. High glycodelin in the medium resulted in signal changes of genes that are part of hypercytokinemia/hyperchemokinemia in the pathogenesis of Influenza and the neuroinflammation signaling pathway. Furthermore, a cell-to-cell interaction pathway between dendritic cells and natural killer cells and a cell-cycle related pathway were affected.

Selected genes of interest identified from IPA analyses were validated by qPCR (Supplemental Fig. S3E). In both immune cell lines, *TNF* and *CXCL10* were most strongly upregulated genes after 24 h of glycodelin stimulation but also Nuclear factor kappa-light-chain-enhancer of activated B cells (*NF-KB)* signaling appeared to be upregulated at earlier treatment time points.

### Spatial analysis of glycodelin and leukocyte markers in NSCLC tissue

The *in vitro* experiments revealed that glycodelin secreted by NSCLC cells is capable of binding to immune cells and affects inflammation related pathways in these cells. To validate these findings *in vivo*, we used multiplex immunofluorescence stainings. In a first step, FFPE samples of the patients were examined from whom the NSCLC cell lines 4950T and 170162T with known glycodelin expression were derived (Fig. 4A). In these sections, glycodelin staining was mainly detected in the tumor areas but also in surrounding stroma. Here, distinct cells were detected being double positive for glycodelin and cluster of differentiation (CD45, white arrows).

**Fig. 4.**
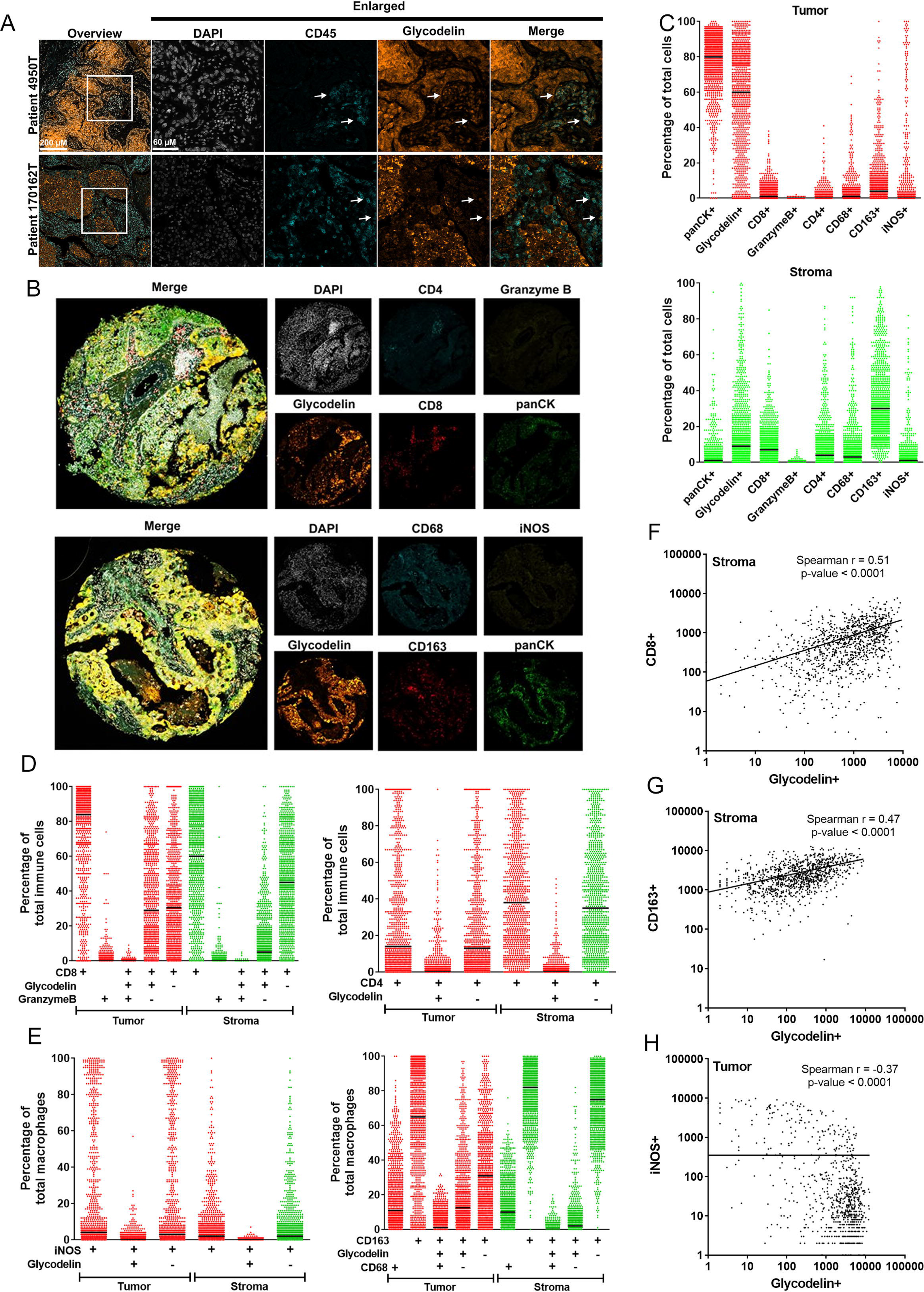
Spatial analyses of glycodelin binding to immune cells *in vivo*. A) Colocalization studies of CD45+ cells and glycodelin in the tumors of 4950T and 170162T. White arrows indicate colocalizing signals. B) Representative stainings for glycodelin and a T cell marker panel as well as a macrophage panel. C) Percentage distribution of the different cell types in tumor and stroma. D) Percentage of single, double and triple positive cells in tumor and stroma for glycodelin and the indicated T cell markers. E) Percentage of single, double and triple positive cells in tumor and stroma for glycodelin and the indicated macrophage markers. F)-H) Spearman correlation analyses of glycodelin and CD8+ cells (F), CD163+ cells (G) and iNOS+ cells (H) in tumor and stroma, respectively. P-value < 0.05 was considered significant. R > 0.5 was considered as correlation.

To gain comprehensive insights into the properties of glycodelin in NSCLC regarding the interaction with immune cells, spatial analysis was performed on tissue microarrays (TMAs) covering 726 patients. The TMAs were stained with two different 5-plex panels (Fig. 4B). In line with the previous findings, glycodelin staining in tumor and stroma was heterogenous (Fig. 4C). pan cytokeratin (CK)+ cells were mainly found in the tumors while immune cells were predominantly detected in stroma. Most of the glycodelin signal found in the tumor tissue was overlapping with panCK staining (Supplemental Fig. S4A and B). We observed a strong association between glycodelin and CD8+ cells but not CD4+ cells, neither in the tumor nor in stroma (Fig. 4D). The number of Granzyme B positive cells was too low for additional analyses. A large proportion of tissue punches was positive for the M1 macrophage marker neuronal nitric oxide synthases (iNOS) in the tumor region, whereas the majority of CD163+ M2 macrophages were situated in the stroma (Fig. 4E). CD68+ macrophages were detected in both areas. Cells triple positive for CD163, glycodelin, and CD68 were detected significantly less than cells that were only positive for CD163 and glycodelin. Cells double positive for glycodelin and iNOS were rarely detected.

In a Spearman correlation analyses we observed a medium correlation (r=0.51) between the CD8+ and glycodelin+ cells in the stroma and a trend (r=0.4) in the tumors of the same patients (Fig. 4F and Supplemental Fig. S4C). The combination of glycodelin and CD163 in the stroma showed a positive correlation (r=0.47, Fig. 4G) at a similar level also to CD68+ cells in the stroma (Supplemental Fig. S4J). In contrast, cells positive for iNOS or glycodelin revealed a weakly negative correlation in the tumor with r=-0.37 (Fig. 4H). None of the other markers correlated with glycodelin (Supplemental Fig. S4D-K).

Based on the multi-immunofluorescence stainings, we investigated the influence of the detection of glycodelin and the co-localization with immune cells on the overall survival of the patients (Fig. 5). Higher glycodelin levels in the stroma (Supplemental Fig. S5A) or in the tumor (Supplemental Fig. S5B) area were generally correlated with a worse overall survival prognosis. Both, male and female patients with a higher staining of glycodelin in the stroma showed a worse overall survival prognosis (Fig. 5A and B). In the tumor area, glycodelin staining was prognostic only for the survival of female patients (Fig. 5C) while the glycodelin staining in male patients (Fig. 5D) showed no differences in relation to overall survival. We also investigated the overall survival prognosis based on our findings of a potential binding of glycodelin to M1 and M2 macrophages. While the co-localization of glycodelin and the M1 macrophage marker iNOS in the stroma correlated with in a significantly better survival prognosis (Fig. 5E), the opposite effect was found for the co-localization of glycodelin and CD163, a M2 macrophage marker (Fig. 5F). The analyses in the tumor area showed similar results (Supplemental Fig. S5C and D). However, generally, the correlation of the staining with the overall survival prognosis was higher in the tumor stroma compared to tumor itself. The potential binding of glycodelin to CD8+ cells was connected with a worse prognosis (Supplemental Fig. S5E and F).

**Fig. 5:**
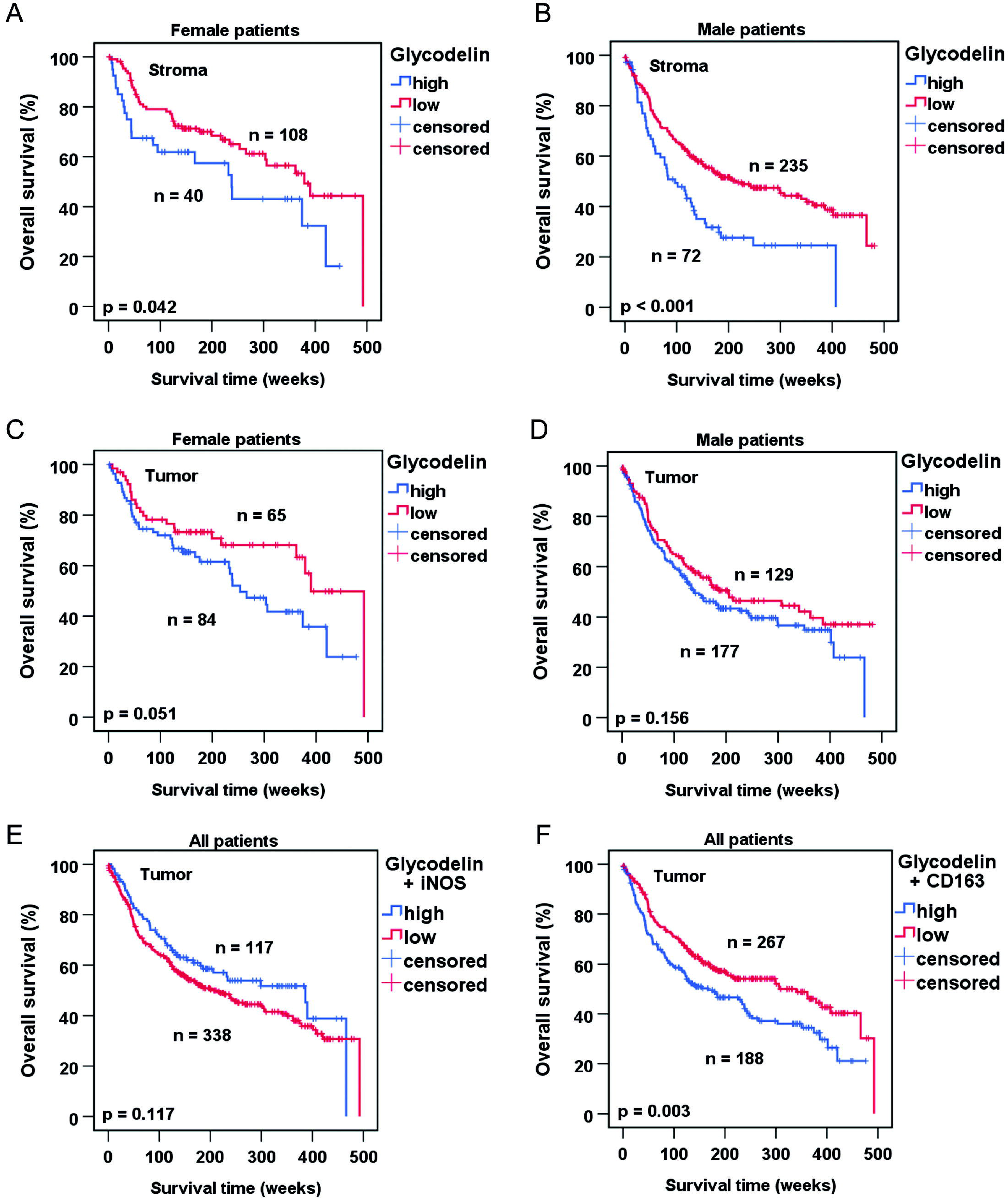
Influence of glycodelin and the colocalization to immune cells on overall survival of patients. A-F. Overall survival analyses based on the multi IF stainings from Fig. 4. Cut-offs for Kaplan-Meier plots were calculated using the Cutoff-Finder (17). p<0.05 was considered significant.

### Glycodelin serum levels predict the clinical benefit of PD-1/PD-L1 therapy in female patients

Immunotherapy for inoperable patients (stage IIIB/IV) with NSCLC is a novel approach that has led to improved treatment outcomes and prolonged progression-free and overall survival in many patients. However, several patients do not benefit from this costly treatment option (2).

As we postulate that glycodelin regulates the tumor microenvironment by binding to immune cells (22), we investigated whether glycodelin serum levels correlate with progression-free survival (PFS) upon immunotherapy. All patients in the study were diagnosed with inoperable NSCLC at the time point of investigation and were treated with anti-PD-1 or anti-PD-L1 antibodies (Table 1).

We observed that the glycodelin serum concentrations varies widely before, ranging from 0 to nearly 300 ng/ml. Survival analyses revealed that glycodelin serum concentrations above the median are associated with significantly shortened PFS in all patients (p=0.048, Fig. 6A). However, this observation was not observed in male patients but only in female patients, in whom elevated glycodelin levels were associated with a significantly shorter PFS (p<0.001, Fig. 6B and C). Several studies have shown that hormones, such as progesterone, regulate glycodelin expression (23, 24). To investigate a potential correlation of these hormones with glycodelin expression in patients with NSCLC, the same patient samples were analyzed for serum concentrations of progesterone, testosterone, estrogen and hCG (Supplemental Figure S6A). hCG was measurable in only 39 out of the 125 patients including some with elevated levels up to 11 pg/ml. Testosterone was the only hormone that had no measured levels outside the physiological range with regards to gender-specific expression. Spearman correlation analysis did not reveal any relation between glycodelin and the investigated hormones (Supplemental Fig. S6B). Progesterone was the only hormone that was associated with poor outcome when elevated in the serum of patients with NSCLC (Supplemental Fig. S6C). This observation was restricted to male patients only (Fig. 6D) whereas no progesterone-level dependent difference in PFS was found in female patients (Fig. 6E).

**Figure 6:**
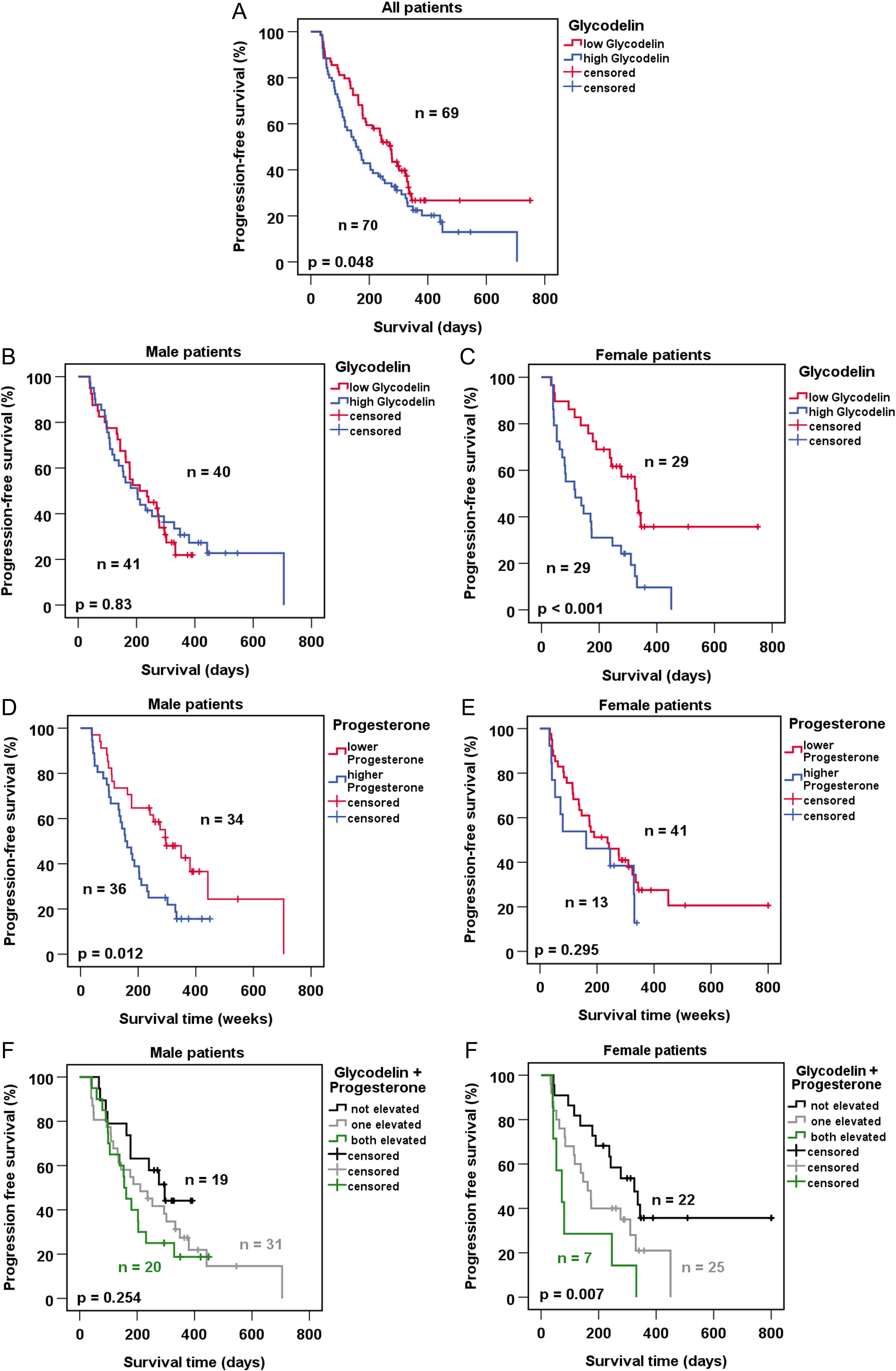
Associations of glycodelin and progesterone serum levels with progression-free survival (PFS) of patients receiving immunotherapy. A) – C) Kaplan-Meier plots using median glycodelin serum levels to divide patients groups. D) and E) Kaplan-Meier plots using median progesterone serum levels to divide patients in two groups. F) and G) Kaplan-Meier plots combining results from measured glycodelin and progesterone serum levels. Patients were devided in a group with both marker values below median, one marker elevated or both values above the median. P < 0.05 was considered significant.

In multivariate Cox regression analysis (Supplemental Table S2) controlling glycodelin and relevant clinical parameters, progesterone turned out to be the only independent prognostic factor for PFS (p=0.026) with an increased hazard ratio (HR=1.694, CI 1.066-2.692) in the investigated cohort. A combination of glycodelin and progesterone serum levels as a 2-marker-panel was not suitable for male patients (Fig. 6F) but significant for PFS of female patients (p=0.007, Fig. 6G). Nevertheless, the combined panel of both markers failed to exceed glycodelin’s potential as biomarker alone (p<0.001, Fig. 6C).

## Discussion

Lung cancer treatment has undergone enormous improvements after the first applications of monoclonal antibodies that target the PD-1/PD-L1 axis (25–27). Nevertheless, overall survival for patients diagnosed at advanced stages remains poor, and there is an unmet clinical need for effective biomarkers and new targets. To address this, we studied glycodelin, which is a reproduction-related, immunosuppressive glycoprotein, expressed also in lung cancer (13, 28–30).

The functionality of the major pregnancy-associated glycodelin glycoform is dependent on its N-linked glycans. Our results revealed significant similarities between the glycans of glycodelin A and NSCLC-derived glycodelin. Furthermore, glycodelin secreted by 4950T cancer cells was sialylated. Such sialylation has been reported to be a driver of immunomodulatory functions of glycodelin A (22, 31–33). Therefore, our results lead to the assumption that glycodelin in NSCLC can potentially act in a similar way as glycodelin A. However, we investigated only glycodelin from two patient-derived cell lines. In future, this finding needs to be validated using patient samples.

Our binding studies showed a glycosylation-independent binding of glycodelin to immune cells. Moreover, preincubation of glycodelin-containing supernatants with an inhibitory antibody resulted in a highly decreased binding to immune cells. Several leukocyte specific receptors have shown to directly bind or modulate binding of pregnancy-related glycodelin, i.e. CD45, CD7, Siglec-6, or L-selectin (34). However, to identify specific receptor(s) was not in the focus of our study since we were able to block the binding to different immune cells efficiently by a glycodelin-specific antibody.

We further examined whether the interaction of NSCLC-derived glycodelin with immune cells is connected to a functional response. Significant genetic changes were detected in THP1 after 3, 8, and 24 h and in KHYG-1 after 24 h, between the cells incubated with media containing high or low levels of glycodelin. The glycodelin regulated genes were related to inflammatory responses and tumor microenvironment pathway. Within these, it is noteworthy that the expression of major regulators like *TNF* and *CXCL10* was increased. The role of TNF interplay between tumor and immune system is not clear since both pro- and antitumorigenic functions have been described for it (35). Interestingly, CXCL10 is also related to INF signaling since TNF is an inducer of CXCL10. CXCL10 has been shown to induce proliferation of tumor cells (36). On the other hand, recruitment of cytotoxic T cells after CXCL10 release has been observed (37). Taken together, previously published results, the fact that glycodelin is highly overexpressed in lung tumors and that this glycodelin-isoform interacts functionally with immune cells suggest a protumorigenic role for glycodelin in lung cancer.

The evaluation of the special analyses of the TMAs indicated a high proportion of glycodelin-positive CD8+ cells in the tumor and the total number of glycodelin positive cells correlated with the number of CD8+ T cells in the stroma. Since activated CD8+ T cells have effector functions and are the main targets of immune checkpoint inhibitors to reactivate an anti-tumor cytotoxic CD8+ T lymphocyte response (38), it is possible that glycodelin suppresses the cytotoxic activity of checkpoint inhibitors. The macrophage panel consisted of CD68 as a general macrophage marker, iNOS as an M1 macrophage marker, and CD163 representing M2 macrophages. Glycodelin and CD163 double positive cells were found in the tumor and stroma region, while a correlation between glycodelin and CD163 was observed in tumor stroma only. In contrast, cell densities of glycodelin positive and iNOS positive macrophages showed a tendency of a negative correlation in tumor sites. While M1 macrophages have been shown to be highly important in the recognition and destruction of cancer cells, M2 macrophages are considered to support tumor growth and metastasis (39). The results indicate an interaction of glycodelin with macrophages. The interaction may drive macrophages towards a M2-like state, and thus, modulate the tumor environment towards an anti-inflammatory and pro-tumorigenic surrounding.

The survival analyses based on the spatial profiles confirmed previous data we have published on RNA level (13). In general, a higher glycodelin protein expression in tumor and stroma tissue resulted in a worse overall survival prognosis for the patients. However, in the tumor, the worse prognosis is restricted to female patients only as already observed before (13). Moreover, the potential binding of glycodelin to immune cells led to a shorter overall survival for the patients when glycodelin is co-localized to CD8+ cells and M2 macrophages. In contrast, binding to M1 macrophages is beneficial for the survival prognosis of the patients. Though, we rarely found iNOS positive cells in the samples supporting the hypothesis that glycodelin secretion from the tumor cells might reorganize the tumor microenvironment to form a pro-tumorigenic niche.

Immune checkpoint inhibition (ICI) represents a promising treatment option for patients with advanced NSCLC (40). The stimulation of coinhibitory pathways modulates the strength and duration of T cell mediated immune responses and prevents damage due to hyperinflammation. These regulatory pathways are exploited by cancer cells to overcome immune surveillance (41). In cell culture experiments an association between glycodelin and PD-L1 expression has already been shown (13). To study the association of glycodelin in therapeutic response to ICI, we measured glycodelin serum levels in patients with stage IIIB/IV NSCLC. We found that elevated glycodelin concentrations are associated with poor outcome but only in female patients. Sex differences in drug-response have been reported regarding the benefit of female patients when treated with epidermal growth factor receptor (EGFR) tyrosine kinase inhibitors (TKIs) (42). With regard to ICI, contradictory results have been reported (43, 44). Additional factors that influence a therapeutic response also include patient characteristics like ethnicity, body mass index (BMI) or the disease-context (45). Investigating differences between female smokers and never-smokers, we observed an increased glycodelin expression in the group of female smokers (46). Consideration of sex as a factor in the application of best possible treatment is still rare in current practice and may need to be considered more in future clinical trials (47, 48). During pregnancy, the expression of the glycoprotein is regulated by different hormones, including progesterone and hCG (7, 24). None of the hormones was correlated with glycodelin concentrations but progesterone levels were negatively associated with the PFS of the male patients. Progesterone interacts with specific progesterone receptors. Studies referring to the receptor functions in the lung (cancer) show contrary results and need to be further investigated (49, 50). At the tumor site, locally elevated amounts of specific hormones, cytokines or other regulators might influence the function of glycodelin and lead to the significant deterioration of ICI treatment in female patients.

Taken together, glycodelin in NSCLC interacts with specific subsets of immune cells in NSCLC tumor tissue and the surrounding stroma. We demonstrated that glycodelin might serve as an easily accessible independent biomarker to predict progression-free survival especially for female patients and may functionally adapt treatment options. Future experiments are needed to further characterize the interaction of glycodelin and patient-derived immune cells and evaluate the potential of glycodelin-inhibitory antibodies alone or in combination with immune checkpoint inhibitors as a treatment option for patients with NSCLC.

## Abbreviations

NSCLC: non-small cell lung cancer;
PAEP: progesterone-associated endometrial protein;
FBS: fetal bovine serum;
siRNA: small interfering ribonucleic acid;
qPCR: quantitative real time polymerase chain reaction;
TMA: tissue multi array,
FFPE: formalin-fixed paraffin-embedded;
hCG: human chorionic gonadotropin;
PFS: progression-free survival;
TTF1: thyroid transcription factor 1;
CK7: cytokeratin 7;
SN: supernatants;
SNA: *Sambucus Nigra* agglutinin;
ECL: *Erythrina cristagalli* lectin;
Con-A: Concanavalin-A;
PSA: *Pisum sativum* agglutinin;
LCS: *Lens culinaris* agglutinin;
TNF: tumor necrosis factor;
CXCL10: C-X-C motif chemokine ligand 10;
CCL4: chemokine (C-C motif) ligand 4;
ICAM1: intercellular adhesion molecule 1;
THBS1: thrombospondin 1;
INFG: interferon gamma;
CCNE2: cell-cycle induction associated genes Cyclin E2;
CDC6: Cell division control protein 6 homolog;
IPA: Ingenuity Pathway Analyses;
TREM1: Triggering receptor expressed on myeloid cells 1;
*NF-K*B: Nuclear factor kappa-light-chain-enhancer of activated B cells;
CD: cluster of differentiation;
CK: cytokeratin;
iNOS: neuronal nitric oxide synthases;
ICI: immune checkpoint inhibition;
EGFR TKI: epidermal growth factor receptor tyrosine kinase inhibitor;
BMI: body mass index

## Acknowledgements

We would like to thank Christa Stolp, Andrea Kreß, Carmen Hoppstock, Martin Fallenbuechel, Elizabeth Xu Meister, and Annikki Ljöfhelm for expert technical support. We would like to thank the NCT tissue bank and Christiane Zorzgelski for providing the TMAs. Moreover, we thank Felix Gruenewald from ROCHE for provision of the glycodelin antibody. All authors have read the journal’s authorship agreement and the manuscript has been reviewed by and approved by all authors. The authors declare that they have no competing interests.

This work was supported by the German Ministry of Education and Research (BMBF), as part of the German Center for Lung Research (82DZL00402, 82DZL001A5, 82DZL00404). HK acknowledges support from Sigrid Jusélius Foundation.

## Availability of data and materials

Affymetrix gene expression data have been deposited in NCBI’s Gene Expression Omnibus and are accessible through GEO Series accession number GSE222124. Original immunoblot images are accessible on request.

