## Supplementary material for "The pregnancy-associated protein glycodelin as a potential sex-specific target for resistance to immunotherapy in non-small cell lung cancer": Supplemental figure legends.docx

**Figure S1: Characterization of 170162T cells and 4950T cells by immunofluorescence analyses.** Both cell lines were stained for adenocarcinoma specific markers cytokeratin 7 (CK7) and thyroid transcription factor 1 (TTF1).

**Figure S2: Binding of glycodelin to cancer immune cell lines.** A) Jurkat cells, THP1 cells and KHYG-1 cells were incubated with FBS containing medium, with 4950T cell specific medium and with 4950T cell supernatants containing glycodelin for 24 h. Afterwards, cell lysates were used for immunoblot analyses.

**Figure S3: Differentially expressed genes and pathway of THP1 cells and KHYG-1 cells treated with glycodelin.** A) Effect of *PAEP* siRNA-treatment on viability of 4950T cells using increasing *PAEP* siRNA concentrations. B) Supernatants of 4950T cells treated with scrambled siRNA (control) or *PAEP* siRNAs used for THP1 cell and KHYG-1 cell incubation. Supernatants derived from 4950T cells were concentrated using a 30 kDa cutoff filter. C) Viability of immune cells after cultivation with 4950T-derived cell supernatants for 24 h. D) Volcano plots of differentially expressed genes in both cell lines after treatment of glycodelin. Cells were incubated with glycodelin-containing or depleted supernatant from 4950T cells for the indicated time. Exemplary genes are highlighted in red if upregulated and in blue if downregulated. E) Gene expression changes of indicated genes after treatment of THP1 cells (D) or KHYG-1 cells (E) with 4950T cell supernatants were determined by qPCR with three biological replicates each.

**Figure S4. Multiplex IF and correlation analyses.** A) and B) Percentage of single and double positive cells in tumor and stroma for glycodelin and panCK in the T cell and macrophage panel. C-K) Spearman correlation analyses of glycodelin and indicated immune cell markers. P-value < 0.05 was considered significant. R > 0.5 was considered as correlation.

**Figure S5. Overall survival analyses.** A-F. Additional overall survival analyses based on the multi IF stainings from Fig. 4. Cut-offs for Kaplan-Meier plots were calculated using the Cutoff-Finder (20). p<0.05 was considered significant.

**Figure S6: Analyses of hormone serum levels.** A) Measured serum levels of the indicated hormones in the immunotherapy cohort (Table 1). B) Correlation between glycodelin serum concentrations and the indicated hormones. C) Progression-free survival of patients receiving immunotherapy in dependency of progesterone. Median serum concentration was used to separate the two groups.
