## Supplementary material for "The pregnancy-associated protein glycodelin as a potential sex-specific target for resistance to immunotherapy in non-small cell lung cancer": Tabel S1.docx

**Table S1. Most regulated genes in immune cells**

| **Most regulated genes in THP1 after 3h** | | | | | | |
| --- | --- | --- | --- | --- | --- | --- |
| ID | Glycodelin Avg (log2) | no Glycodelin Avg (log2) | Fold Change | P-val | FDR P-val | Gene Symbol |
| 204103_at | 7,76 | 4,48 | 9,71 | 1,01E-05 | 0,009 | CCL4 |
| 204533_at | 7,43 | 4,33 | 8,56 | 4,84E-07 | 0,0013 | CXCL10 |
| 202859_x_at | 8,01 | 4,92 | 8,52 | 1,31E-08 | 0,0001 | CXCL8 |
| 202643_s_at | 9,11 | 6,37 | 6,68 | 5,48E-09 | 9,98E-05 | TNFAIP3 |
| 202644_s_at | 10,58 | 7,9 | 6,43 | 1,44E-08 | 0,0001 | TNFAIP3 |
| 206765_at | 6,35 | 3,75 | 6,06 | 6,89E-07 | 0,0015 | KCNJ2 |
| 221477_s_at | 9,66 | 7,13 | 5,77 | 1,52E-08 | 0,0001 | LOC100129518; SOD2 |
| 211506_s_at | 6,54 | 4,03 | 5,66 | 2,07E-05 | 0,013 | CXCL8 |
| 1566342_at | 10,14 | 7,65 | 5,63 | 5,15E-09 | 9,98E-05 | SOD2 |
| 232504_at | 5,78 | 3,33 | 5,49 | 6,95E-07 | 0,0015 | LOC285628 |
| 205114_s_at | 8,36 | 5,93 | 5,4 | 6,69E-07 | 0,0015 | CCL3; CCL3L1; CCL3L3 |
| 207113_s_at | 10,08 | 7,73 | 5,09 | 2,04E-05 | 0,0129 | TNF |
| 206026_s_at | 5,29 | 2,95 | 5,04 | 6,50E-07 | 0,0015 | TNFAIP6 |
| 206157_at | 7,22 | 4,89 | 5 | 3,87E-06 | 0,0044 | PTX3 |
| 202638_s_at | 6,89 | 4,59 | 4,95 | 4,64E-09 | 9,98E-05 | ICAM1 |
| 238727_at | 5,87 | 3,73 | 4,4 | 0,0001 | 0,0393 |  |
| 231779_at | 8,6 | 6,5 | 4,31 | 4,78E-07 | 0,0013 | IRAK2 |
| 231513_at | 5,5 | 3,43 | 4,2 | 5,01E-07 | 0,0013 |  |
| 204897_at | 10,82 | 8,75 | 4,18 | 4,11E-08 | 0,0002 | PTGER4 |
| 210538_s_at | 5,45 | 3,4 | 4,16 | 7,11E-08 | 0,0003 | BIRC3 |
| 236982_at | 6,93 | 4,89 | 4,12 | 1,05E-06 | 0,0018 |  |
| 206025_s_at | 5,66 | 3,63 | 4,07 | 2,59E-05 | 0,0154 | TNFAIP6 |
| 210260_s_at | 9,12 | 7,25 | 3,66 | 3,68E-07 | 0,0012 | TNFAIP8 |
| 215223_s_at | 9,43 | 7,63 | 3,48 | 1,58E-07 | 0,0006 | LOC100129518; SOD2 |
| 209288_s_at | 9,52 | 7,77 | 3,38 | 1,76E-08 | 0,0001 | CDC42EP3 |
| 222162_s_at | 7,23 | 5,51 | 3,3 | 2,42E-06 | 0,0035 | ADAMTS1 |
| 239459_s_at | 5,74 | 4,03 | 3,27 | 0,0013 | 0,1794 | CYP19A1 |
| 208296_x_at | 8,75 | 7,04 | 3,25 | 3,37E-06 | 0,0041 | TNFAIP8 |
| 223218_s_at | 8,09 | 6,42 | 3,2 | 4,11E-08 | 0,0002 | NFKBIZ |
| 244414_at | 6,69 | 5,03 | 3,17 | 8,85E-05 | 0,0345 |  |
| 230966_at | 6,65 | 4,99 | 3,14 | 0,0012 | 0,173 | IL4I1 |
| 216841_s_at | 8,65 | 7,01 | 3,12 | 1,07E-06 | 0,0018 | LOC100129518; SOD2 |
| 209286_at | 8,43 | 6,78 | 3,12 | 2,92E-06 | 0,004 | CDC42EP3 |
| 222486_s_at | 5,54 | 3,92 | 3,08 | 3,11E-08 | 0,0002 | ADAMTS1 |
| 239876_at | 7,44 | 5,85 | 3,01 | 0,0007 | 0,1278 |  |
| 1558569_at | 6,43 | 4,85 | 3 | 0,0001 | 0,0419 | LOC100131541 |
| 220091_at | 7,73 | 6,2 | 2,9 | 1,58E-05 | 0,0114 | SLC2A6 |
| 225685_at | 8,78 | 7,27 | 2,86 | 2,54E-06 | 0,0036 | CDC42EP3 |
| 227481_at | 5,97 | 4,45 | 2,86 | 0,0002 | 0,0497 | CNKSR3 |
| 242405_at | 6,43 | 4,94 | 2,8 | 0,0002 | 0,0597 |  |
| 219908_at | 6,67 | 5,21 | 2,75 | 8,47E-06 | 0,0078 | DKK2 |
| 239649_at | 6,77 | 5,31 | 2,74 | 0,0019 | 0,2195 |  |
| 209774_x_at | 5,47 | 4,02 | 2,73 | 1,81E-05 | 0,0122 | CXCL2 |
| 239460_at | 6,92 | 5,47 | 2,73 | 0,0041 | 0,3377 | CYP19A1 |
| 204470_at | 5,39 | 3,95 | 2,73 | 0,0002 | 0,059 | CXCL1 |
| 201236_s_at | 9,16 | 7,78 | 2,61 | 3,42E-05 | 0,0185 | BTG2 |
| 217767_at | 9,03 | 7,65 | 2,6 | 0,0031 | 0,2857 | C3 |
| 219951_s_at | 10,16 | 8,8 | 2,57 | 3,20E-08 | 0,0002 | DZANK1 |
| 236738_at | 8,04 | 6,7 | 2,55 | 3,90E-05 | 0,0201 | C3orf80 |
| 205681_at | 7,33 | 5,99 | 2,54 | 0,0036 | 0,3128 | BCL2A1 |
| 220252_x_at | 9,12 | 7,77 | 2,53 | 3,37E-06 | 0,0041 | CXorf21 |
| 240863_at | 4,29 | 2,95 | 2,52 | 0,0051 | 0,3739 | CYP19A1 |
| 209892_at | 10,44 | 9,11 | 2,5 | 9,45E-07 | 0,0018 | FUT4 |
| 219721_at | 6,46 | 5,15 | 2,48 | 0,003 | 0,2806 |  |
| 202637_s_at | 6,88 | 5,57 | 2,47 | 2,74E-08 | 0,0002 | ICAM1 |
| 240061_at | 5,7 | 4,42 | 2,43 | 0,0014 | 0,1871 |  |
| 207339_s_at | 7,52 | 6,23 | 2,43 | 0,0049 | 0,3692 | LTB |
| 218810_at | 8,58 | 7,32 | 2,4 | 1,87E-07 | 0,0007 | ZC3H12A |
| 230614_at | 6,54 | 5,29 | 2,37 | 0,0017 | 0,2057 |  |
| 227180_at | 5,35 | 4,1 | 2,37 | 0,005 | 0,3704 | ELOVL7 |
| 209239_at | 9,38 | 8,14 | 2,36 | 2,02E-07 | 0,0007 | NFKB1 |
| 205463_s_at | 8,89 | 7,66 | 2,34 | 0,0023 | 0,2439 | PDGFA |
| 209906_at | 9,31 | 8,09 | 2,34 | 0,0007 | 0,1247 | C3AR1 |
| 204823_at | 6,43 | 5,22 | 2,32 | 0,0055 | 0,394 | NAV3 |
| 207535_s_at | 6,66 | 5,44 | 2,32 | 1,41E-05 | 0,0107 | NFKB2 |
| 203548_s_at | 8,88 | 7,68 | 2,3 | 0,0004 | 0,0888 | LPL |
| 203549_s_at | 8,46 | 7,27 | 2,29 | 0,0004 | 0,086 | LPL |
| 242827_x_at | 6,57 | 5,38 | 2,27 | 0,0029 | 0,272 |  |
| 243808_at | 7,18 | 6 | 2,26 | 0,0063 | 0,4222 |  |
| 209287_s_at | 7,77 | 6,6 | 2,25 | 4,58E-07 | 0,0013 | CDC42EP3 |
| 215078_at | 3,96 | 2,79 | 2,24 | 0,0008 | 0,1419 | LOC100129518; SOD2 |
| 204363_at | 10,11 | 8,94 | 2,24 | 0,0287 | 0,7261 | F3 |
| 228343_at | 6,58 | 5,42 | 2,24 | 0,0009 | 0,1419 | POU2F2 |
| 237981_at | 6,21 | 5,06 | 2,22 | 5,25E-05 | 0,0243 | CMYA5 |
| 205242_at | 4,35 | 3,2 | 2,22 | 0,0008 | 0,1365 | CXCL13 |
| 209946_at | 5,51 | 4,36 | 2,21 | 0,0007 | 0,1279 | VEGFC |
| 203475_at | 7,88 | 6,73 | 2,21 | 0,0027 | 0,2628 | CYP19A1 |
| 204440_at | 8,15 | 7,02 | 2,19 | 4,56E-06 | 0,0051 | CD83 |
| 1562475_at | 4,96 | 3,86 | 2,15 | 4,68E-05 | 0,0228 | TEX41 |
| 204224_s_at | 8,14 | 7,04 | 2,14 | 9,47E-07 | 0,0018 | GCH1 |
| 242907_at | 4,07 | 2,98 | 2,12 | 1,39E-05 | 0,0107 | GBP2 |
| 226302_at | 5,79 | 4,71 | 2,12 | 0,0181 | 0,642 | ATP8B1 |
| 1570505_at | 5,33 | 4,25 | 2,11 | 0,0013 | 0,1834 | ABCB4 |
| 216867_s_at | 8,54 | 7,48 | 2,1 | 0,0051 | 0,3739 | PDGFA |
| 209893_s_at | 8,67 | 7,61 | 2,09 | 1,01E-06 | 0,0018 | FUT4 |
| 229437_at | 8,16 | 7,11 | 2,07 | 1,69E-05 | 0,0116 | MIR155HG |
| 211122_s_at | 3,5 | 2,46 | 2,05 | 0,0001 | 0,047 | CXCL11 |
| 222859_s_at | 6,82 | 5,78 | 2,05 | 0,0022 | 0,2365 | DAPP1 |
| 213943_at | 10,35 | 9,32 | 2,05 | 3,21E-06 | 0,0041 | TWIST1 |
| 228153_at | 4,78 | 3,76 | 2,04 | 0,001 | 0,1512 | RNF144B |
| 218627_at | 7,82 | 6,79 | 2,03 | 0,0005 | 0,109 | DRAM1 |
| 229830_at | 8,65 | 7,63 | 2,02 | 0,0008 | 0,1414 |  |
| 239529_at | 8,96 | 7,95 | 2,02 | 0,0048 | 0,3656 | DCANP1; TIFAB |
| 221143_at | 6,38 | 5,37 | 2,02 | 0,0014 | 0,1911 | RPA4 |
| 216060_s_at | 7,79 | 6,78 | 2,01 | 0,0006 | 0,1145 | DAAM1 |
| 212906_at | 7,74 | 8,74 | -2,01 | 5,98E-07 | 0,0015 | GRAMD1B |
| 202254_at | 5,12 | 6,15 | -2,05 | 0,0001 | 0,0378 | SIPA1L1 |
| 210512_s_at | 9,81 | 10,84 | -2,05 | 0,007 | 0,4447 | VEGFA |
| 213258_at | 4,95 | 6,03 | -2,11 | 3,20E-05 | 0,0176 | TFPI |
| 203761_at | 7,06 | 8,14 | -2,11 | 0,0012 | 0,1774 | SLA |
| 208078_s_at | 6,1 | 7,18 | -2,12 | 2,27E-05 | 0,0139 | LOC102724428; SIK1 |
| 209967_s_at | 6,95 | 8,05 | -2,14 | 5,99E-06 | 0,0064 | CREM |
| 1563357_at | 2,86 | 3,97 | -2,15 | 4,67E-05 | 0,0228 | TNF |
| 200796_s_at | 7,44 | 8,56 | -2,17 | 0,0138 | 0,5906 | MCL1 |
| 210643_at | 4,98 | 6,1 | -2,18 | 0,0129 | 0,5724 | TNFSF11 |
| 212977_at | 4,63 | 5,76 | -2,18 | 0,0005 | 0,1048 | ACKR3 |
| 204912_at | 7,03 | 8,17 | -2,21 | 6,30E-06 | 0,0066 | IL10RA |
| 230707_at | 5,48 | 6,64 | -2,23 | 0,0014 | 0,1839 | SORL1 |
| 202948_at | 6 | 7,19 | -2,28 | 0,0203 | 0,6683 | IL1R1 |
| 206337_at | 4,36 | 5,56 | -2,29 | 0,0015 | 0,1927 | CCR7 |
| 227646_at | 3,65 | 4,85 | -2,3 | 0,0002 | 0,057 | EBF1 |
| 228499_at | 7,51 | 8,73 | -2,33 | 1,92E-06 | 0,0031 | PFKFB4 |
| 217738_at | 7,22 | 8,44 | -2,34 | 1,12E-05 | 0,0095 | NAMPT |
| 211527_x_at | 5,85 | 7,09 | -2,37 | 0,0027 | 0,267 | VEGFA |
| 210772_at | 4,36 | 5,6 | -2,37 | 0,0002 | 0,0497 | FPR2 |
| 236220_at | 3,78 | 5,04 | -2,39 | 5,38E-05 | 0,0247 | SLC16A10 |
| 203887_s_at | 10,55 | 11,81 | -2,4 | 2,85E-05 | 0,0164 | THBD |
| 209723_at | 7,54 | 8,81 | -2,41 | 0,0001 | 0,0449 | SERPINB9 |
| 203888_at | 9,95 | 11,23 | -2,43 | 1,36E-05 | 0,0107 | THBD |
| 217996_at | 5,91 | 7,21 | -2,45 | 0,0394 | 0,7817 | PHLDA1 |
| 226844_at | 4,18 | 5,5 | -2,49 | 2,33E-06 | 0,0034 | MOB3B |
| 229568_at | 3,68 | 5,03 | -2,55 | 2,41E-05 | 0,0145 | MOB3B |
| 201041_s_at | 6,91 | 8,26 | -2,55 | 8,84E-07 | 0,0018 | DUSP1 |
| 220330_s_at | 4,49 | 5,88 | -2,64 | 0,0001 | 0,0421 | SAMSN1 |
| 205119_s_at | 6,09 | 7,54 | -2,74 | 7,80E-05 | 0,0316 | FPR1 |
| 206877_at | 4,73 | 6,2 | -2,77 | 0,0002 | 0,0632 | MXD1 |
| 237252_at | 7,83 | 9,31 | -2,79 | 3,10E-06 | 0,004 | THBD |
| 221009_s_at | 6,36 | 7,84 | -2,8 | 0,0009 | 0,1419 | ANGPTL4 |
| 225842_at | 6,24 | 7,73 | -2,82 | 0,0015 | 0,1947 | PHLDA1 |
| 226275_at | 7,43 | 8,94 | -2,85 | 8,12E-06 | 0,0078 | MXD1 |
| 206118_at | 5,62 | 7,27 | -3,14 | 8,51E-05 | 0,0334 | STAT4 |
| 228846_at | 7,52 | 9,18 | -3,15 | 6,93E-05 | 0,0295 | MXD1 |
| 223333_s_at | 5,07 | 6,94 | -3,65 | 0,001 | 0,1585 | ANGPTL4 |
| 201108_s_at | 4,64 | 6,72 | -4,23 | 1,03E-05 | 0,009 | THBS1 |
| 228186_s_at | 4,61 | 7,14 | -5,78 | 2,21E-06 | 0,0034 | RSPO3 |
| 205239_at | 6,3 | 9,02 | -6,57 | 0,0035 | 0,3086 | AREG |
| 201109_s_at | 4,87 | 7,89 | -8,1 | 2,28E-06 | 0,0034 | THBS1 |
| 201110_s_at | 5,31 | 8,82 | -11,35 | 1,00E-06 | 0,0018 | THBS1 |
| **Most regulated genes in THP1 after 8h** | | | | | | |
| ID | Glycodelin Avg (log2) | no Glycodelin Avg (log2) | Fold Change | P-val | FDR P-val | Gene Symbol |
| 215223_s_at | 9,67 | 7,53 | 4,42 | 1,14E-10 | 6,24E-06 | LOC100129518; SOD2 |
| 205943_at | 7,05 | 5 | 4,14 | 1,54E-07 | 0,0014 | TDO2 |
| 204533_at | 6,68 | 4,77 | 3,76 | 5,04E-07 | 0,0023 | CXCL10 |
| 1566342_at | 9,5 | 7,61 | 3,72 | 2,22E-09 | 6,06E-05 | SOD2 |
| 221477_s_at | 8,68 | 6,8 | 3,67 | 3,63E-07 | 0,0022 | LOC100129518; SOD2 |
| 207113_s_at | 8,14 | 6,27 | 3,65 | 0,0011 | 0,1675 | TNF |
| 204103_at | 6,26 | 4,54 | 3,3 | 2,37E-05 | 0,0202 | CCL4 |
| 219424_at | 5,83 | 4,12 | 3,27 | 1,55E-06 | 0,0039 | EBI3 |
| 230966_at | 6,75 | 5,04 | 3,25 | 5,28E-05 | 0,0331 | IL4I1 |
| 205242_at | 7,4 | 5,72 | 3,21 | 4,55E-05 | 0,0311 | CXCL13 |
| 216841_s_at | 8,6 | 7,01 | 3 | 1,74E-08 | 0,0003 | LOC100129518; SOD2 |
| 220091_at | 7,67 | 6,09 | 2,98 | 3,04E-08 | 0,0004 | SLC2A6 |
| 204475_at | 4,68 | 3,12 | 2,95 | 0,0008 | 0,1382 | MMP1 |
| 202638_s_at | 6,23 | 4,69 | 2,92 | 3,50E-07 | 0,0022 | ICAM1 |
| 203562_at | 5,94 | 4,46 | 2,78 | 0,0003 | 0,0817 | FEZ1 |
| 221159_at | 5,39 | 3,97 | 2,68 | 3,22E-05 | 0,0241 |  |
| 205114_s_at | 7,22 | 5,81 | 2,66 | 0,003 | 0,2712 | CCL3; CCL3L1; CCL3L3 |
| 212190_at | 9,63 | 8,26 | 2,59 | 1,45E-06 | 0,0039 | SERPINE2 |
| 239459_s_at | 5,58 | 4,23 | 2,55 | 0,0022 | 0,2351 | CYP19A1 |
| 211597_s_at | 5,39 | 4,07 | 2,51 | 5,82E-08 | 0,0006 | HOPX |
| 232504_at | 4,91 | 3,6 | 2,48 | 1,02E-06 | 0,0031 | LOC285628 |
| 220146_at | 4,8 | 3,51 | 2,45 | 0,0029 | 0,2652 | TLR7 |
| 207819_s_at | 6,66 | 5,37 | 2,44 | 1,10E-05 | 0,0127 | ABCB4 |
| 209875_s_at | 8,54 | 7,27 | 2,41 | 0,02 | 0,578 | SPP1 |
| 209946_at | 5,32 | 4,08 | 2,37 | 0,0012 | 0,1738 | VEGFC |
| 230614_at | 6,44 | 5,22 | 2,32 | 0,0031 | 0,2719 |  |
| 202644_s_at | 9,69 | 8,51 | 2,27 | 3,02E-06 | 0,0059 | TNFAIP3 |
| 211122_s_at | 4,02 | 2,84 | 2,27 | 4,55E-05 | 0,0311 | CXCL11 |
| 229830_at | 6,96 | 5,81 | 2,23 | 0,0003 | 0,0801 |  |
| 205463_s_at | 7,71 | 6,56 | 2,22 | 0,0002 | 0,0649 | PDGFA |
| 231513_at | 7,49 | 6,33 | 2,22 | 1,56E-06 | 0,0039 |  |
| 209994_s_at | 6,43 | 5,29 | 2,21 | 5,92E-05 | 0,0349 | ABCB1; ABCB4 |
| 219721_at | 6,51 | 5,37 | 2,2 | 0,0055 | 0,3591 |  |
| 216867_s_at | 7,88 | 6,74 | 2,2 | 0,0004 | 0,0886 | PDGFA |
| 228060_at | 7,35 | 6,22 | 2,19 | 8,01E-07 | 0,0031 | SLC35F1 |
| 219938_s_at | 7,74 | 6,62 | 2,18 | 0,0001 | 0,0551 | PSTPIP2 |
| 202643_s_at | 8,31 | 7,19 | 2,18 | 4,78E-07 | 0,0023 | TNFAIP3 |
| 203475_at | 9,18 | 8,06 | 2,17 | 1,98E-06 | 0,0044 | CYP19A1 |
| 207339_s_at | 6,81 | 5,7 | 2,16 | 0,0021 | 0,2292 | LTB |
| 205404_at | 9,57 | 8,46 | 2,16 | 2,05E-07 | 0,0016 | HSD11B1 |
| 239460_at | 6,62 | 5,53 | 2,14 | 0,0102 | 0,4683 | CYP19A1 |
| 236471_at | 4,24 | 3,17 | 2,1 | 0,0036 | 0,2919 | NFE2L3 |
| 206765_at | 8,15 | 7,09 | 2,09 | 6,20E-05 | 0,0349 | KCNJ2 |
| 228153_at | 5,02 | 3,98 | 2,06 | 0,0014 | 0,1874 | RNF144B |
| 202748_at | 6,05 | 5,03 | 2,02 | 3,97E-05 | 0,0285 | GBP2 |
| 237032_x_at | 4,89 | 5,93 | -2,05 | 3,16E-05 | 0,024 | SIPA1L1 |
| 227948_at | 4,67 | 5,71 | -2,06 | 3,64E-06 | 0,0066 | FGD4 |
| 229584_at | 4,79 | 5,87 | -2,11 | 8,64E-05 | 0,0435 | LRRK2 |
| 214841_at | 4,44 | 5,52 | -2,12 | 0,0013 | 0,1864 | CNIH3 |
| 227052_at | 5,45 | 6,57 | -2,16 | 5,72E-06 | 0,0087 | SMIM14 |
| 211959_at | 6,51 | 7,63 | -2,18 | 0,0013 | 0,1864 | IGFBP5 |
| 224894_at | 4,59 | 5,82 | -2,35 | 0,0002 | 0,0728 | YAP1 |
| 210772_at | 4,73 | 6,09 | -2,55 | 6,10E-05 | 0,0349 | FPR2 |
| 226834_at | 4,42 | 5,79 | -2,59 | 0,0028 | 0,2613 | CLMP |
| 201109_s_at | 3,86 | 5,24 | -2,6 | 0,0171 | 0,5461 | THBS1 |
| 205119_s_at | 7,17 | 8,79 | -3,06 | 0,0002 | 0,0654 | FPR1 |
| 227662_at | 4,61 | 6,24 | -3,09 | 0,0008 | 0,1449 | SYNPO2 |
| 210664_s_at | 5,15 | 6,97 | -3,51 | 9,55E-07 | 0,0031 | TFPI |
| 201110_s_at | 3,88 | 5,71 | -3,54 | 0,0076 | 0,4158 | THBS1 |
| 213258_at | 5,06 | 6,99 | -3,83 | 1,16E-06 | 0,0033 | TFPI |
| 205239_at | 5,33 | 7,88 | -5,88 | 0,002 | 0,227 | AREG |
| **Most regulated genes in THP1 after 24h** | | | | | | |
| ID | Glycodelin Avg (log2) | no Glycodelin Avg (log2) | Fold Change | P-val | FDR P-val | Gene Symbol |
| 211122_s_at | 10,66 | 7,56 | 8,6 | 1,32E-06 | 0,0021 | CXCL11 |
| 210163_at | 8,15 | 5,25 | 7,47 | 1,75E-06 | 0,0027 | CXCL11 |
| 231702_at | 7,39 | 4,51 | 7,39 | 4,77E-07 | 0,0017 | TDO2 |
| 204103_at | 9,02 | 6,3 | 6,58 | 0,0002 | 0,086 | CCL4 |
| 216598_s_at | 10,57 | 7,86 | 6,55 | 2,76E-08 | 0,0002 | CCL2 |
| 205943_at | 11,46 | 8,78 | 6,38 | 2,95E-07 | 0,0011 | TDO2 |
| 219424_at | 7,54 | 4,91 | 6,17 | 8,08E-07 | 0,0019 | EBI3 |
| 205114_s_at | 10,03 | 7,67 | 5,13 | 0,0008 | 0,2019 | CCL3; CCL3L1; CCL3L3 |
| 204475_at | 7,58 | 5,37 | 4,62 | 8,90E-11 | 4,86E-06 | MMP1 |
| 204533_at | 12,94 | 10,74 | 4,59 | 8,21E-07 | 0,0019 | CXCL10 |
| 203131_at | 8,92 | 6,76 | 4,46 | 5,42E-07 | 0,0017 | PDGFRA |
| 214038_at | 11,52 | 9,43 | 4,27 | 6,36E-06 | 0,0074 | CCL8 |
| 206134_at | 8,13 | 6,12 | 4,04 | 1,04E-06 | 0,0019 | ADAMDEC1 |
| 230966_at | 8,66 | 6,68 | 3,93 | 6,62E-05 | 0,0397 | IL4I1 |
| 215223_s_at | 10,49 | 8,51 | 3,93 | 1,73E-08 | 0,0002 | LOC100129518; SOD2 |
| 206026_s_at | 5,7 | 3,76 | 3,83 | 2,39E-05 | 0,0182 | TNFAIP6 |
| 207113_s_at | 8,37 | 6,44 | 3,81 | 0,0046 | 0,5024 | TNF |
| 203936_s_at | 9,86 | 7,96 | 3,74 | 5,78E-08 | 0,0004 | MMP9 |
| 202638_s_at | 7,75 | 5,88 | 3,66 | 8,70E-07 | 0,0019 | ICAM1 |
| 202859_x_at | 8,42 | 6,59 | 3,55 | 0,003 | 0,4061 | CXCL8 |
| 212143_s_at | 10,46 | 8,68 | 3,45 | 8,70E-05 | 0,0485 | IGFBP3 |
| 221477_s_at | 9,61 | 7,83 | 3,42 | 5,27E-06 | 0,0063 | LOC100129518; SOD2 |
| 220146_at | 6,71 | 4,97 | 3,33 | 7,29E-07 | 0,0019 | TLR7 |
| 229560_at | 6,94 | 5,25 | 3,21 | 3,09E-08 | 0,0002 | TLR8 |
| 1566342_at | 9,96 | 8,28 | 3,19 | 2,76E-05 | 0,0204 | SOD2 |
| 218810_at | 8,79 | 7,14 | 3,14 | 5,37E-07 | 0,0017 | ZC3H12A |
| 232504_at | 5,47 | 3,82 | 3,14 | 9,34E-07 | 0,0019 | LOC285628 |
| 203868_s_at | 5,24 | 3,59 | 3,13 | 8,43E-08 | 0,0005 | VCAM1 |
| 229435_at | 5,68 | 4,06 | 3,07 | 0,0001 | 0,0618 | GLIS3 |
| 205242_at | 10,13 | 8,53 | 3,04 | 4,74E-06 | 0,0058 | CXCL13 |
| 216841_s_at | 9,5 | 7,9 | 3,03 | 2,57E-07 | 0,0011 | LOC100129518; SOD2 |
| 205959_at | 6,21 | 4,65 | 2,95 | 2,70E-07 | 0,0011 | MMP13 |
| 210095_s_at | 11,53 | 9,98 | 2,93 | 8,93E-05 | 0,0488 | IGFBP3 |
| 230258_at | 5,85 | 4,32 | 2,87 | 0,0004 | 0,1254 | GLIS3 |
| 229947_at | 5,76 | 4,25 | 2,84 | 2,25E-06 | 0,0031 | PI15 |
| 205463_s_at | 8,38 | 6,87 | 2,84 | 3,24E-09 | 6,85E-05 | PDGFA |
| 229830_at | 8,13 | 6,65 | 2,78 | 3,76E-09 | 6,85E-05 |  |
| 202637_s_at | 7,8 | 6,33 | 2,76 | 5,66E-07 | 0,0017 | ICAM1 |
| 211506_s_at | 7,07 | 5,62 | 2,72 | 0,0002 | 0,089 | CXCL8 |
| 216834_at | 9,39 | 7,95 | 2,72 | 0,0101 | 0,6765 | RGS1 |
| 212190_at | 10,77 | 9,34 | 2,7 | 1,09E-07 | 0,0006 | SERPINE2 |
| 206157_at | 7,03 | 5,6 | 2,68 | 0,0003 | 0,1083 | PTX3 |
| 211597_s_at | 8 | 6,59 | 2,67 | 1,15E-06 | 0,002 | HOPX |
| 210873_x_at | 5,99 | 4,58 | 2,66 | 0,0001 | 0,0618 | APOBEC3A; APOBEC3A_B |
| 227834_at | 5,19 | 3,78 | 2,65 | 0,0002 | 0,0892 | TXLNB |
| 202643_s_at | 9,11 | 7,7 | 2,65 | 8,98E-07 | 0,0019 | TNFAIP3 |
| 208607_s_at | 6,93 | 5,52 | 2,65 | 2,99E-06 | 0,004 | SAA1; SAA2; SAA2-SAA4 |
| 230233_at | 4,19 | 2,8 | 2,63 | 0,0002 | 0,0735 |  |
| 204961_s_at | 10,32 | 8,95 | 2,58 | 1,83E-05 | 0,0151 | NCF1; NCF1B; NCF1C |
| 220091_at | 8,44 | 7,08 | 2,56 | 1,87E-06 | 0,0028 | SLC2A6 |
| 227487_s_at | 5,49 | 4,16 | 2,52 | 0,001 | 0,2306 | SERPINE2 |
| 202988_s_at | 7,48 | 6,15 | 2,51 | 0,0104 | 0,6818 | RGS1 |
| 210839_s_at | 6,88 | 5,55 | 2,51 | 1,56E-05 | 0,0137 | ENPP2 |
| 201631_s_at | 7,62 | 6,29 | 2,51 | 1,00E-06 | 0,0019 | IER3 |
| 203828_s_at | 6,1 | 4,77 | 2,5 | 1,66E-06 | 0,0026 | IL32 |
| 231577_s_at | 8,49 | 7,16 | 2,5 | 0,0006 | 0,1609 | GBP1 |
| 209670_at | 6,09 | 4,77 | 2,5 | 2,12E-06 | 0,0031 | TRAC |
| 1555756_a_at | 6,21 | 4,89 | 2,49 | 0,0002 | 0,0737 | CLEC7A |
| 205681_at | 7,13 | 5,82 | 2,49 | 0,0006 | 0,1706 | BCL2A1 |
| 203915_at | 6,35 | 5,05 | 2,48 | 1,49E-05 | 0,0136 | CXCL9 |
| 206025_s_at | 5,72 | 4,43 | 2,45 | 2,82E-05 | 0,0205 | TNFAIP6 |
| 238727_at | 5,9 | 4,61 | 2,45 | 0,0007 | 0,1836 |  |
| 211719_x_at | 8,38 | 7,11 | 2,42 | 0,0073 | 0,6075 | FN1 |
| 202270_at | 7,42 | 6,15 | 2,41 | 0,0003 | 0,0943 | GBP1 |
| 204470_at | 6,59 | 5,33 | 2,4 | 0,0004 | 0,1262 | CXCL1 |
| 209774_x_at | 5,61 | 4,35 | 2,4 | 0,0015 | 0,2883 | CXCL2 |
| 223217_s_at | 6,8 | 5,54 | 2,39 | 7,48E-07 | 0,0019 | NFKBIZ |
| 214084_x_at | 10,38 | 9,12 | 2,38 | 1,16E-05 | 0,0116 | NCF1 |
| 221345_at | 5,97 | 4,74 | 2,36 | 0,0014 | 0,2773 | FFAR2 |
| 209684_at | 8,89 | 7,67 | 2,34 | 0,0009 | 0,2141 | RIN2 |
| 216867_s_at | 8,59 | 7,37 | 2,32 | 2,48E-06 | 0,0034 | PDGFA |
| 207536_s_at | 6,11 | 4,89 | 2,32 | 1,37E-08 | 0,0002 | TNFRSF9 |
| 201739_at | 10,16 | 8,95 | 2,31 | 0,0042 | 0,4932 | SGK1 |
| 202269_x_at | 8,58 | 7,39 | 2,28 | 0,0006 | 0,1609 | GBP1 |
| 209392_at | 7,64 | 6,46 | 2,27 | 0,0001 | 0,0681 | ENPP2 |
| 231779_at | 7,61 | 6,44 | 2,25 | 0,0016 | 0,2883 | IRAK2 |
| 223218_s_at | 8,51 | 7,34 | 2,25 | 1,20E-05 | 0,0117 | NFKBIZ |
| 205226_at | 6,88 | 5,72 | 2,24 | 1,99E-05 | 0,016 | PDGFRL |
| 210004_at | 4,65 | 3,49 | 2,24 | 1,04E-06 | 0,0019 | OLR1 |
| 1553141_at | 7,14 | 5,99 | 2,23 | 2,40E-05 | 0,0182 | LACC1 |
| 1552658_a_at | 6,24 | 5,09 | 2,23 | 1,00E-05 | 0,0102 | NAV3 |
| 229327_s_at | 5,89 | 4,75 | 2,21 | 0,0011 | 0,2366 |  |
| 217546_at | 4,76 | 3,62 | 2,2 | 0,0003 | 0,0901 | MT1M |
| 202644_s_at | 10,27 | 9,14 | 2,18 | 7,47E-06 | 0,0085 | TNFAIP3 |
| 210258_at | 5,28 | 4,16 | 2,17 | 0,0039 | 0,4706 | RGS13 |
| 205476_at | 10,21 | 9,09 | 2,16 | 0,0014 | 0,2765 | CCL20 |
| 205067_at | 8,6 | 7,48 | 2,16 | 9,87E-06 | 0,0102 | IL1B |
| 215078_at | 4,53 | 3,42 | 2,16 | 0,0062 | 0,5715 | LOC100129518; SOD2 |
| 219386_s_at | 9,4 | 8,29 | 2,16 | 0,0006 | 0,1718 | SLAMF8 |
| 210495_x_at | 8,65 | 7,54 | 2,16 | 0,0046 | 0,5051 | FN1 |
| 226237_at | 3,78 | 2,67 | 2,16 | 4,05E-06 | 0,0051 | COL8A1 |
| 215990_s_at | 8,18 | 7,08 | 2,15 | 0,0024 | 0,3448 | BCL6 |
| 216442_x_at | 8,69 | 7,6 | 2,14 | 0,0063 | 0,5722 | FN1 |
| 227458_at | 5,92 | 4,83 | 2,13 | 0,0007 | 0,1792 | CD274 |
| 202833_s_at | 8,76 | 7,68 | 2,11 | 9,12E-06 | 0,0101 | SERPINA1 |
| 203477_at | 9,07 | 8 | 2,11 | 2,52E-05 | 0,0189 | COL15A1 |
| 209031_at | 7 | 5,93 | 2,1 | 9,73E-07 | 0,0019 | CADM1 |
| 212464_s_at | 8,53 | 7,46 | 2,1 | 0,0059 | 0,5545 | FN1 |
| 208025_s_at | 5,59 | 4,54 | 2,08 | 0,0103 | 0,6818 | HMGA2 |
| 202672_s_at | 7,94 | 6,91 | 2,04 | 0,0023 | 0,3419 | ATF3 |
| 236471_at | 5,29 | 4,26 | 2,04 | 0,0002 | 0,0727 | NFE2L3 |
| 231513_at | 7,87 | 6,85 | 2,04 | 7,93E-07 | 0,0019 |  |
| 1555759_a_at | 10,38 | 9,36 | 2,03 | 0,0002 | 0,0694 | CCL5 |
| 206513_at | 7,56 | 6,55 | 2,03 | 0,0004 | 0,1347 | AIM2 |
| 39402_at | 8,51 | 7,49 | 2,02 | 5,52E-05 | 0,0355 | IL1B |
| 220832_at | 5,09 | 4,09 | 2 | 2,85E-05 | 0,0205 | TLR8 |
| 209699_x_at | 7,55 | 8,6 | -2,06 | 0,0205 | 0,8096 | AKR1C2 |
| **Most regulated genes in KHYG-1 after 24h** | | | | | | |
| ID | Glycodelin Avg (log2) | no Glycodelin Avg (log2) | Fold Change | P-val | FDR P-val | Gene Symbol |
| 202330_s_at | 6,17 | 4,36 | 3,52 | 0,0382 | 0,3908 | UNG |
| 205034_at | 5,73 | 3,93 | 3,49 | 0,0178 | 0,3905 | CCNE2 |
| 211814_s_at | 5,31 | 3,57 | 3,33 | 0,0405 | 0,3923 | CCNE2 |
| 229450_at | 4,97 | 3,25 | 3,3 | 0,0017 | 0,3905 | IFIT3 |
| 210354_at | 5,1 | 3,44 | 3,17 | 0,0002 | 0,3905 | IFNG |
| 204103_at | 11,04 | 9,38 | 3,16 | 0,0003 | 0,3905 | CCL4 |
| 216248_s_at | 4,36 | 2,78 | 2,99 | 4,09E-05 | 0,3905 | NR4A2 |
| 203967_at | 6,61 | 5,1 | 2,85 | 0,034 | 0,3908 | CDC6 |
| 220651_s_at | 4,89 | 3,4 | 2,81 | 0,0054 | 0,3905 | MCM10 |
| 203126_at | 5,55 | 4,06 | 2,8 | 0,0158 | 0,3905 | IMPA2 |
| 225655_at | 7,63 | 6,16 | 2,77 | 0,0237 | 0,3905 | UHRF1 |
| 204622_x_at | 4,92 | 3,45 | 2,76 | 5,40E-05 | 0,3905 | NR4A2 |
| 206365_at | 7,15 | 5,69 | 2,76 | 0,0001 | 0,3905 | XCL1 |
| 207535_s_at | 5,67 | 4,24 | 2,68 | 0,0114 | 0,3905 | NFKB2 |
| 1557129_a_at | 3,99 | 2,61 | 2,6 | 0,0013 | 0,3905 | FAM111B |
| 219288_at | 5,57 | 4,21 | 2,56 | 0,0028 | 0,3905 | C3orf14 |
| 228033_at | 5,17 | 3,84 | 2,5 | 0,0134 | 0,3905 | E2F7 |
| 221521_s_at | 7,33 | 6,03 | 2,46 | 0,033 | 0,3908 | GINS2 |
| 219990_at | 6,41 | 5,15 | 2,39 | 0,0112 | 0,3905 | E2F8 |
| 209891_at | 6,91 | 5,67 | 2,37 | 0,0014 | 0,3905 | SPC25 |
| 1552470_a_at | 4,57 | 3,34 | 2,35 | 0,0191 | 0,3905 | ABHD11 |
| 210567_s_at | 5,47 | 4,24 | 2,34 | 0,0034 | 0,3905 | SKP2 |
| 231164_at | 6,72 | 5,5 | 2,33 | 0,0038 | 0,3905 | ABCA17P |
| 211162_x_at | 6,73 | 5,53 | 2,3 | 0,0003 | 0,3905 | SCD |
| 222680_s_at | 7,26 | 6,09 | 2,26 | 0,0308 | 0,3908 | DTL |
| 207113_s_at | 6,63 | 5,46 | 2,25 | 0,0003 | 0,3905 | TNF |
| 204621_s_at | 3,91 | 2,74 | 2,25 | 0,0032 | 0,3905 | NR4A2 |
| 207339_s_at | 9,08 | 7,92 | 2,23 | 0,0162 | 0,3905 | LTB |
| 203210_s_at | 6,54 | 5,41 | 2,18 | 0,0103 | 0,3905 | RFC5 |
| 223307_at | 8,34 | 7,22 | 2,18 | 0,0001 | 0,3905 | CDCA3 |
| 202068_s_at | 7,58 | 6,46 | 2,17 | 0,0041 | 0,3905 | LDLR |
| 242918_at | 5,35 | 4,24 | 2,16 | 0,0029 | 0,3905 | NASP |
| 242069_at | 5,79 | 4,68 | 2,16 | 0,0019 | 0,3905 | CBX5 |
| 212791_at | 5,79 | 4,68 | 2,16 | 0,019 | 0,3905 | C1orf216 |
| 216228_s_at | 4,59 | 3,48 | 2,16 | 0,0054 | 0,3905 | WDHD1 |
