## Supplementary material for "The pregnancy-associated protein glycodelin as a potential sex-specific target for resistance to immunotherapy in non-small cell lung cancer": Tabel S2.docx

**Table S2: Cox-regression of immunotherapy cohort**

CI = confidencial interval, ECOG = Eastern Cooperative Oncology Group

| **Parameter** | **Hazard Ratio (95% CI)** | **p-value** |
| --- | --- | --- |
| Glycodelin high | 1.458 (0.899-2.365) | 0.127 |
| Progesterone high | 1.694 (1.066-2.692) | **0.026** |
| Immunotherapie line | 1.149 (0.876-1.506) | 0.316 |
| Sex (f vs male) | 1.347 (0.849-2.136) | 0.206 |
| Age at diagnosis | 1.016 (0.989-1.043) | 0.251 |
| ECOG | 1.068 (0.683-1.671) | 0.771 |
| Stage at diagnosis | 0.871 (0.659-1.151) | 0.331 |
