## Supplementary material for "The pregnancy-associated protein glycodelin as a potential sex-specific target for resistance to immunotherapy in non-small cell lung cancer": Table 1.docx

T**able 1. Clinical parameters of the investigated patient cohorts**

NOS = not otherwise specified, n.d. = no data, ECOG = Eastern Cooperative Oncology Group, mAb = monoclonal antibody, LUSC = Lung squamous cell carcinoma, LUAD = Lung adenocarcinoma, LCC = Large cell carcinoma, NOS = not otherwise specified, SCC = Small-cell lung cancer, TMA = tissue multi array

| Cohort characteristics - Immunotherapy serum cohort | | | | | | |
| --- | --- | --- | --- | --- | --- | --- |
| **Parameter** | **n** | **(%)** |  | **Parameter** | **n** | **(%)** |
| ***Median Age*** | 63  (38-85) |  |  | ***Line***  ***Immunotherapy*** |  |  |
| Total | **139** |  |  | 1st | 77 | 55 |
|  |  |  |  | 2nd | 53 | 38 |
| ***Gender*** |  |  |  | 3rd | 6 | 4 |
| Male | 81 | 58 |  | 4th | 3 | 2 |
| Female | 58 | 42 |  |  |  |  |
|  |  |  |  | ***mAb therapy*** |  |  |
| ***Histology*** |  |  |  | PD-1 | 114 | 82 |
| LUSC | 31 | 22 |  | PD-L1 | 25 | 18 |
| LUAD | 97 | 70 |  |  |  |  |
| LCC | 3 | 2 |  | ***ECOG*** |  |  |
| NOS | 8 | 6 |  | 0 | 57 | 41 |
|  |  |  |  | 1 | 74 | 53 |
| ***PD-L1*** |  |  |  | 2 | 2 | 1 |
| <1 % | 24 | 17 |  | n.d. | 6 | 4 |
| 1-49 % | 54 | 39 |  |  |  |  |
| >50 % | 45 | 32 |  |  |  |  |
| n.d. | 16 | 12 |  |  |  |  |
| **Cohort characteristics – Multi IF tissue cohort (TMA)** | | | | | | |
| **Parameter** | **n** | **(%)** |  | **Parameter** | **n** | **(%)** |
| ***Median Age*** | 63  (38-86) |  |  | ***ECOG*** |  |  |
| Total | 726 |  |  | 0 | 508 | 70 |
|  |  |  |  | 1 | 191 | 26 |
| ***Gender*** |  |  |  | 2 | 25 | 3 |
| Male | 484 | 67 |  | 3 | 1 | 0 |
| Female | 242 | 33 |  | 4 | 1 | 0 |
| ***Histology*** |  |  |  | ***Pathological stage (7th edition)*** |  |  |
| LUSC | 247 | 34 |  | I | 228 | 31 |
| LUAD | 374 | 52 |  | II | 192 | 26 |
| NSCLC mix | 45 | 6 |  | III | 261 | 36 |
| LCC | 34 | 5 |  | IV | 36 | 5 |
| NOS | 25 | 3 |  | n.d. | 9 | 1 |
| SCC | 1 | 0 |  |  |  |  |
| ***Smoking status*** |  |  |  |  |  |  |
| Current smoker | 209 | 29 |  |  |  |  |
| Ex-smoker | 421 | 58 |  |  |  |  |
| Never smoker | 72 | 10 |  |  |  |  |
| n.d. | 24 | 3 |  |  |  |  |
