## Supplementary figures and images for "The pregnancy-associated protein glycodelin as a potential sex-specific target for resistance to immunotherapy in non-small cell lung cancer"

### Figure S1 4950T and 170162T.jpg

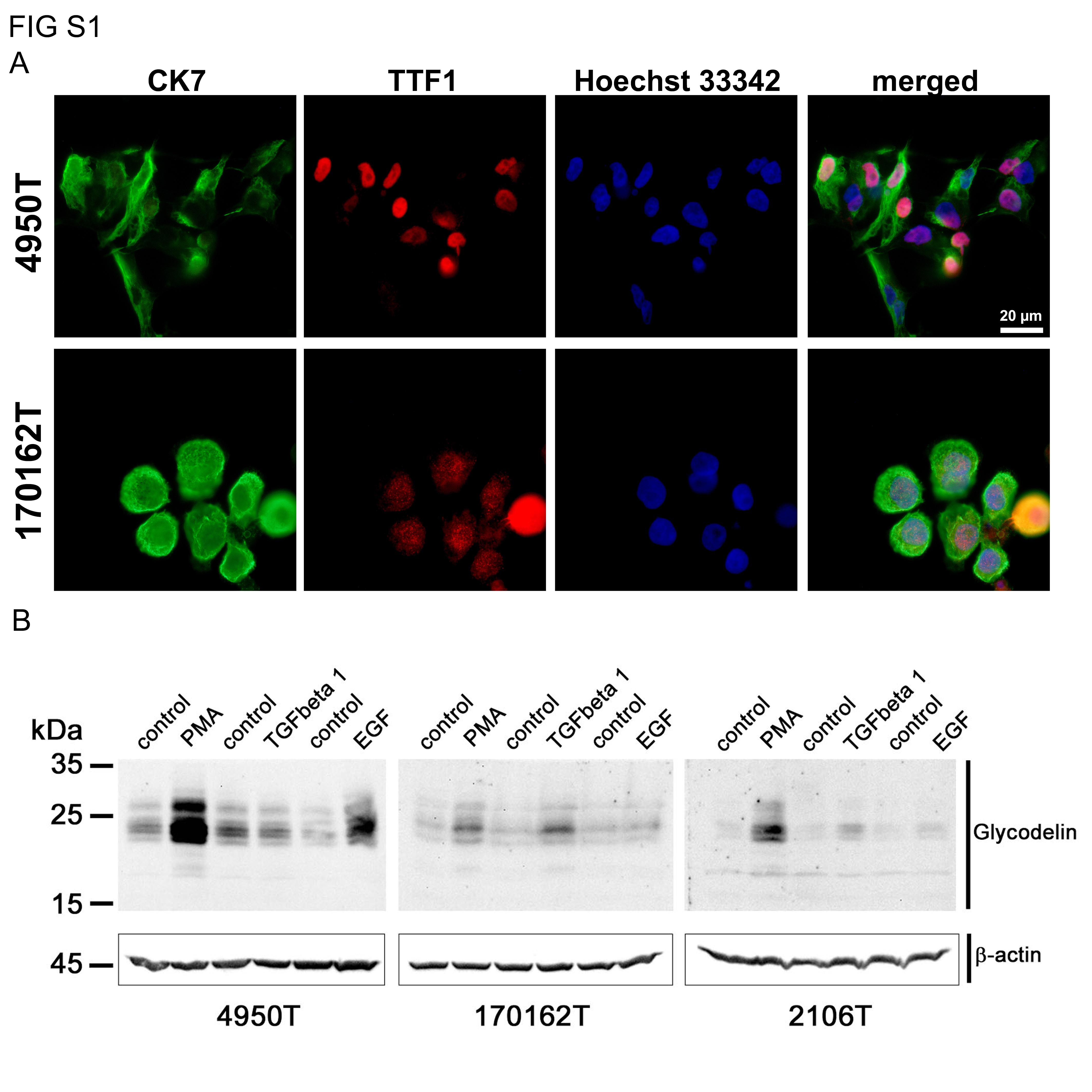

### Figure S2 Binding Assays.jpg

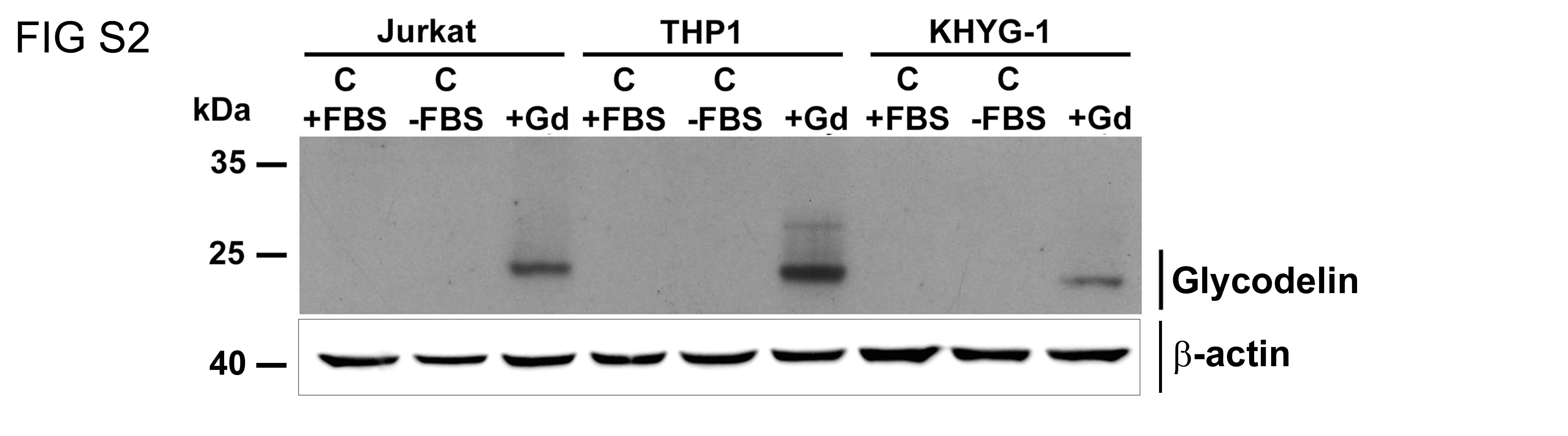

### Figure S3 Affymetrix.jpg

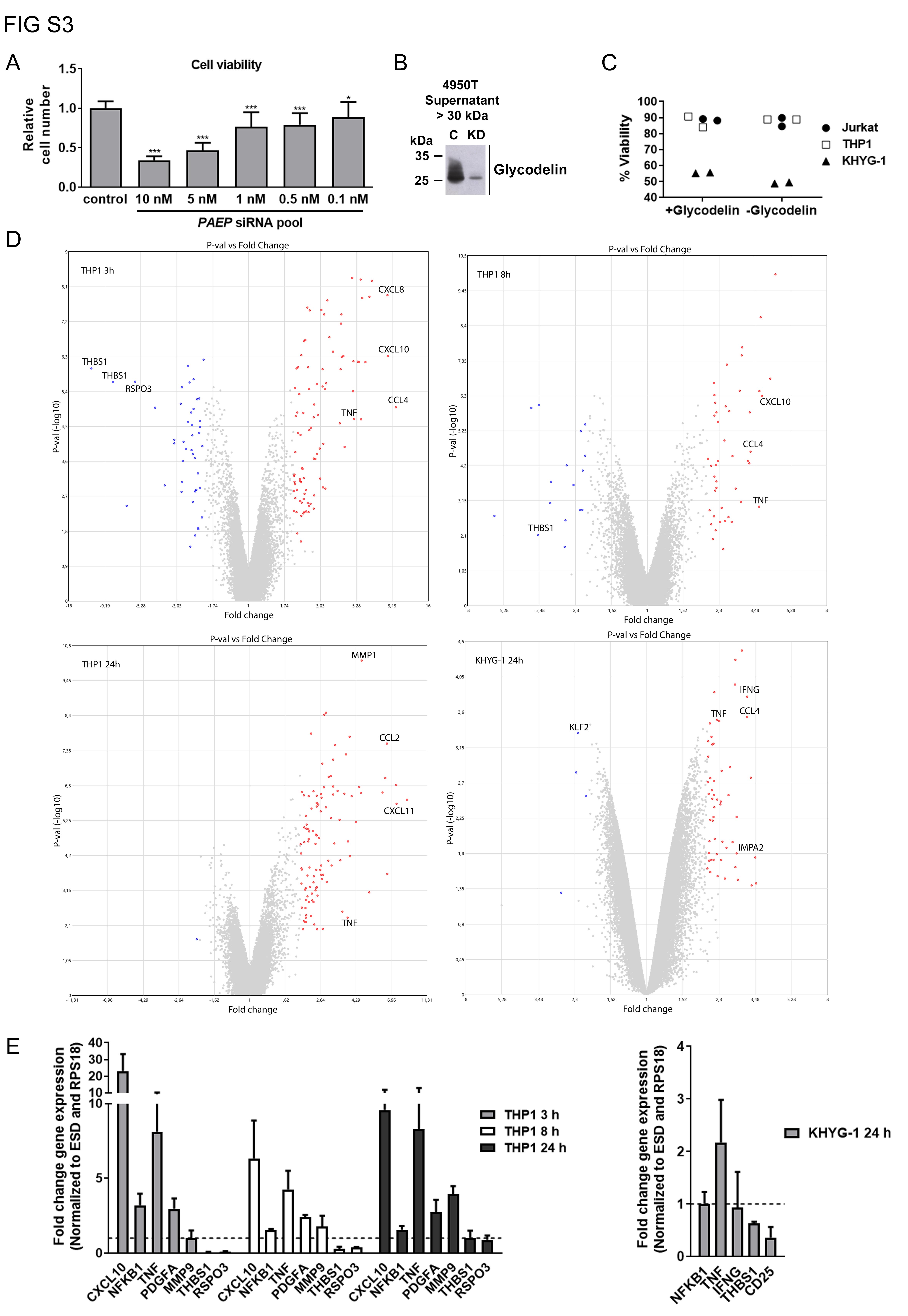

### Figure S4 multiplex IF.jpg

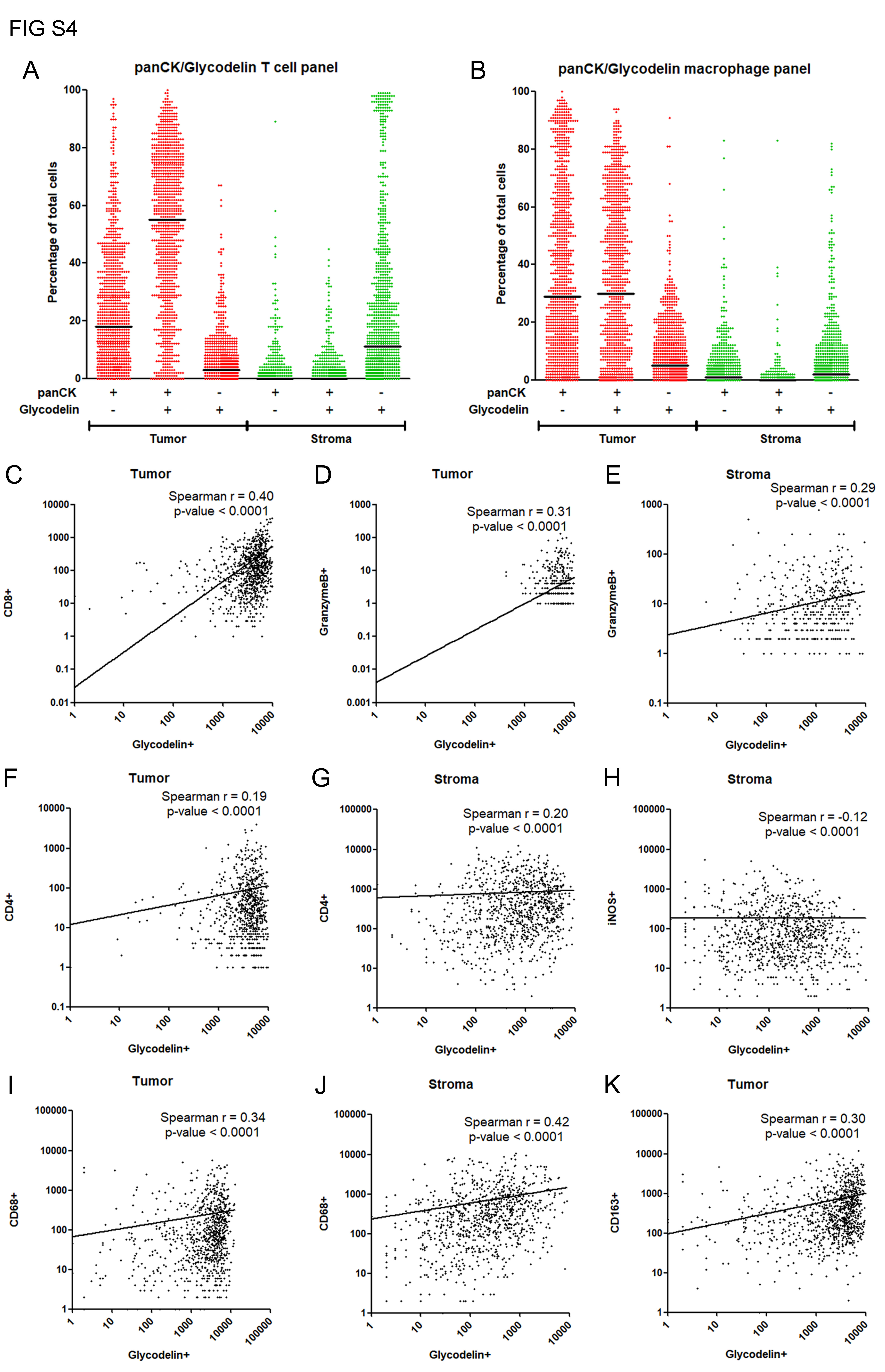

### Figure S5 Survival.jpg

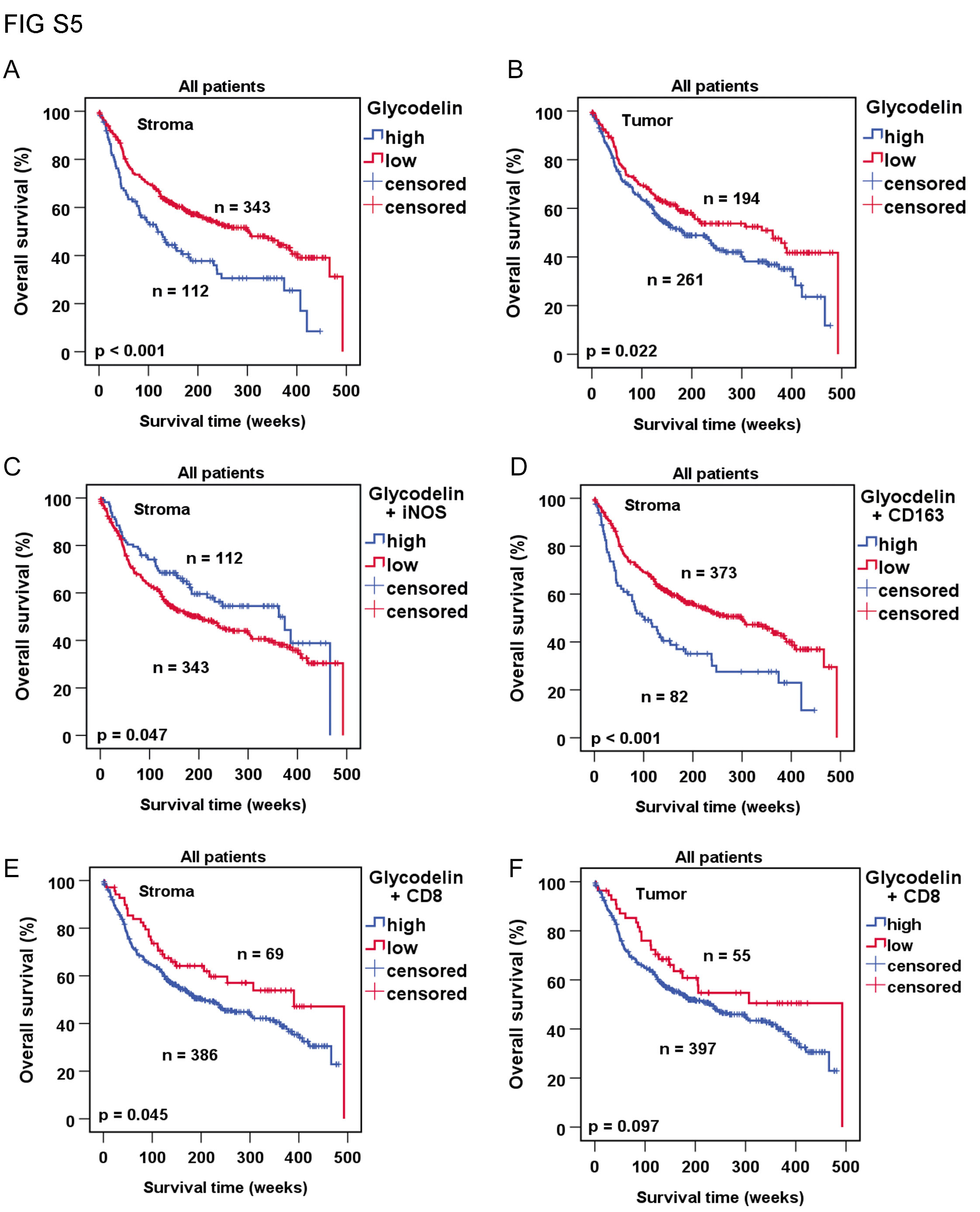

### Figure S6 Hormones.jpg

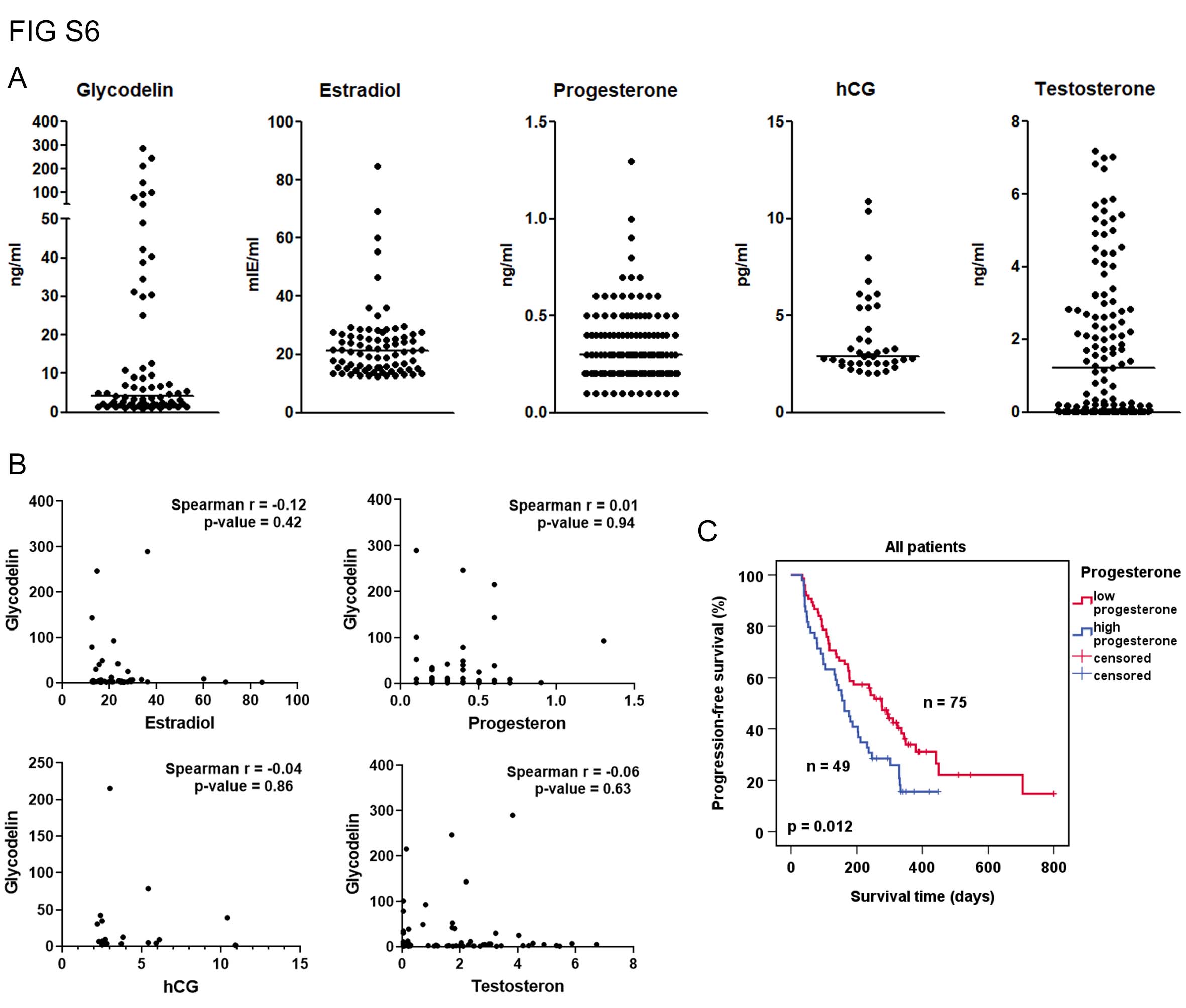
